# Alzheimer’s Disease-Associated Proteomic Changes in Olfactory Stem Cells Show Pathway-Level Transcriptomic Overlap

**DOI:** 10.64898/2026.10.19.760270

**Authors:** Laura M. Rantanen, Harley Robinson, Danielle Dutra Voigt, Xochitl Diaz, Riikka Lampinen, Sonja Mykkänen, Juho Kalapudas, Anne M. Koivisto, Elina Penttilä, Lotta E. Oikari, Hazel Quek, Katja M. Kanninen, Anthony R. White

## Abstract

Alzheimer’s disease (AD) is a progressive, neurodegenerative disorder and the leading cause of dementia worldwide. Although recent therapeutic advances have provided some disease-modifying potential, there is currently no curative treatment for AD, highlighting the need for improved disease models and early biomarkers. Olfactory mucosal stem cells, which can be expanded *in vitro* as olfactory neurosphere-derived (ONS) cells, represent a promising source for biomarker discovery because olfactory tissue is more accessible than central nervous system tissue and has been reported to exhibit AD-related pathological changes. In this study, we used subcellular fractionation-based proteomics, reconstructed bulk proteomics and integrated transcriptomic-proteomic analyses to characterise alterations in ONS cells derived from healthy control, mild cognitive impairment (MCI) and AD individuals. Fraction-specific proteomics identified cytoplasmic and nuclear-enriched protein abundance changes, including alterations in proteins and pathways associated with cytoskeletal organisation, RNA processing, intracellular trafficking, autophagy and mitochondrial/metabolic processes. Reduced LC3B abundance in cytoplasmic AD ONS cells was supported by western blot analysis, although LC3B abundance alone does not indicate altered autophagic flux. Reconstructed bulk proteomics identified 40 FDR-significant proteins in AD compared with HC, defining an AD-associated protein signature enriched for extracellular matrix, cytoskeletal, organelle-associated and RNA-processing pathways. Despite proteomic enrichment of mitochondrial and metabolic pathways, Seahorse Mito Stress Test analysis did not identify significant group-level differences in mitochondrial respiration. Integrated transcriptomic-proteomic analysis revealed partial RNA-protein concordance and pathway-level overlap across disease-associated processes. Together, these findings support the use of patient-derived ONS cells as a model for investigating AD- and MCI-associated molecular changes and highlight protein signatures for further validation.

## 2. Introduction

Alzheimer’s disease (AD) is the leading cause of dementia, contributing to approximately 60-70% of dementia cases globally (World Health Organisation, 2026). It is a progressive and currently irreversible neurodegenerative disease that typically begins with mild memory impairment and advances to severe cognitive decline, functional disability and loss of independence [1]. Around 55 million people worldwide are currently living with dementia [2], and this number is predicted to increase substantially in the coming decades in the absence of effective disease-modifying treatments [3,4]. In 2019, the global economic cost of dementia was estimated at USD 1.3 trillion [5], highlighting the major public health and socioeconomic burden associated with AD and related dementias. The neurodegenerative process underlying AD develops over several decades [6], with cognitive symptoms typically emerging only after years of asymptomatic neuropathological change [7,6]. Mild cognitive impairment (MCI) is often considered a prodromal stage of dementia, although not all individuals with MCI progress to AD or another dementia [8,9,7]. This makes MCI an important stage for investigating early disease-associated molecular changes. The high failure rate of clinical trials for AD therapeutics suggests that disease-modifying interventions may be more effective if applied earlier in the disease process, before widespread neurodegeneration has occurred [6,10]. Therefore, improved understanding of early cellular and molecular changes associated with AD progression, together with the identification of accessible biomarkers, remains a major priority.

Olfaction, the sense of smell, is closely connected with central nervous system function [11,12]. Olfactory sensory neurons located in the olfactory epithelium detect odorants in the nasal cavity and project through the cribriform plate to the olfactory bulb, which is directly connected to brain regions involved in emotions, memory and cognition, including the amygdala, entorhinal cortex and hippocampus [13,14] (**Fig. 1A**). Olfactory dysfunction is commonly reported in neurodegenerative diseases, including AD and Parkinson’s disease, and it is one of the earliest clinical features of AD. Deficits in odour threshold, identification and recognition memory can occur before clear cognitive decline and may reflect early deterioration in brain regions involved in episodic memory and perceptual processing [15,16]. Although the mechanisms underlying olfactory dysfunction in AD are not fully understood, previous studies have linked olfactory impairment with amyloid-β, phosphorylated tau pathology and APOE genotype [15]. Amyloid-β and phosphorylated tau have also been detected in *post-mortem* olfactory epithelium [17,18] and nasal secretions from individuals with AD [19,20] supporting the olfactory system as a potential source of disease-relevant biomarkers.

**Fig. 1.**
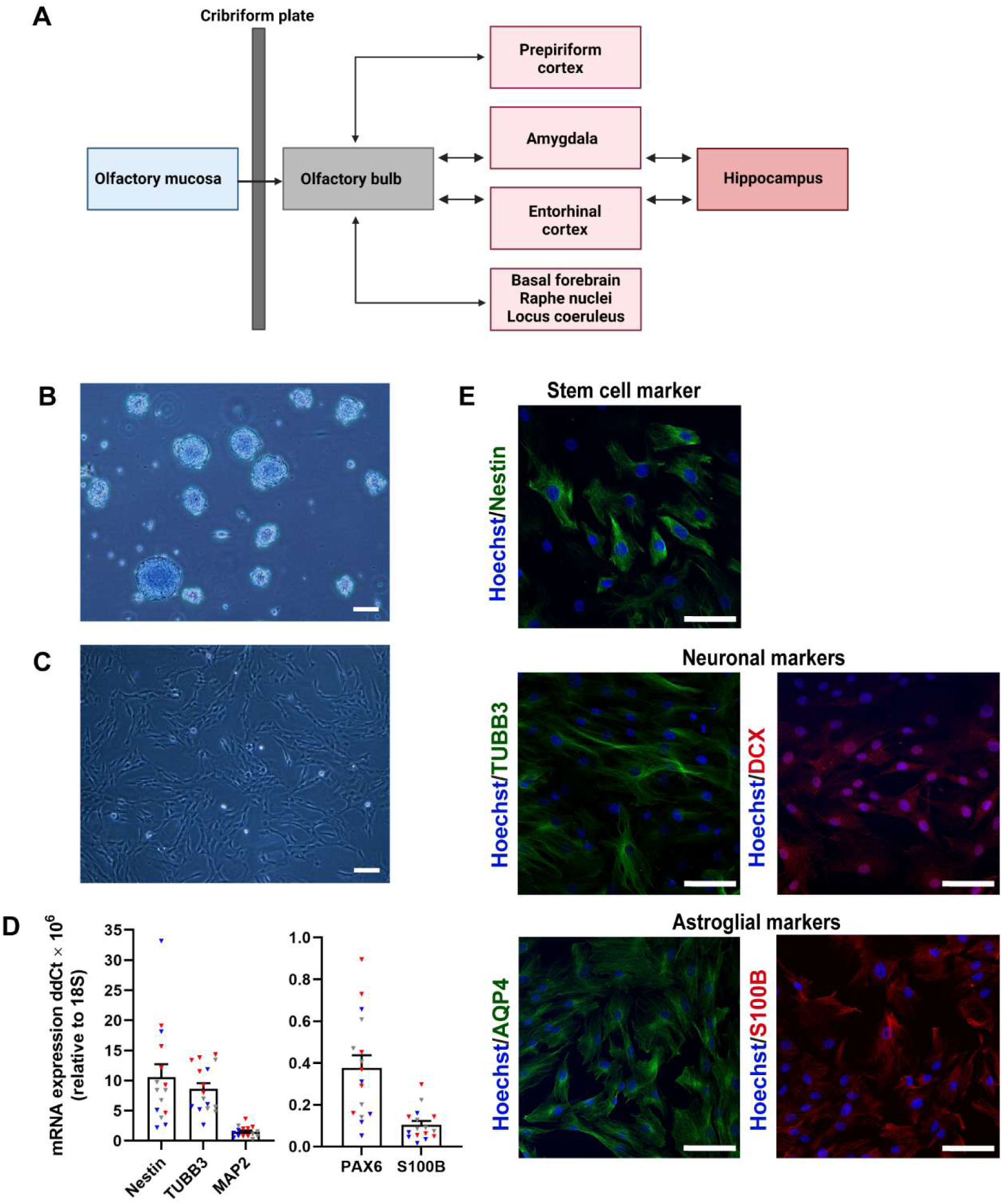
Characterisation of patient-derived olfactory neurosphere-derived cells. (**A**) Schematic showing olfactory processing-related regions known to show early neurodegenerative changes in AD, including the olfactory bulb, prepiriform cortex, amygdala, entorhinal cortex, locus coeruleus and hippocampus. Adapted from [28]. Created with BioRender.com. (**B**) Phenotype of olfactory neurospheres generated from cultured olfactory mucosal cells (4× Magnification). Scale bar, 100 µm. (**C**) Phenotype of olfactory neurosphere-derived (ONS) cells cultured as an adherent monolayer in serum-containing medium (4× Magnification). Scale bar, 100 µm. (**D**) qRT-PCR characterisation of ONS cells showing relative mRNA expression of stem cell-associated markers Nestin and PAX6, immature neuronal markers TUBB3 and MAP2 and the astroglial marker S100B across HC, MCI and AD ONS cell lines. HC *n* = 5 (blue), MCI *n* = 5 (grey) and AD *n* = 6 (red), with three technical replicates per cell line. (**E**) Immunocytochemistry of patient-derived ONS cells showing expression of stem cell markers Nestin (green), neuronal markers TUBB3 (green) and DCX (red), and astroglial markers AQP4 (green) and S100B (red) (20× Magnification). Nuclei were counterstained with Hoechst. Scale bars, 100 µm.

The olfactory mucosa (OM) also contains basal stem and progenitor cells, including horizontal and globose basal cells, which support lifelong regeneration of olfactory neurons and other epithelial cell types [12,21,22]. Patient-derived cells from the olfactory mucosa provide a useful model for investigating AD- and MCI-associated molecular changes, with the broader aim of identifying disease-relevant pathways and candidate biomarker signatures that may support earlier detection and a better understanding of disease progression. As described in our previous study, olfactory mucosal stem cells are accessible through nasal biopsy, can be expanded as olfactory neurosphere-derived cells (ONS cells), and retain disease-relevant molecular features associated with AD and MCI [23]. These cells offer several advantages for biomarker discovery, including accessibility, potential for longitudinal sampling, and biological relevance to olfactory pathways affected early in AD [23,17,24–26].

In the present study, we combined subcellular fractionation proteomics, reconstructed bulk proteomics and transcriptomic-proteomic integration analyses to characterise disease-associated molecular changes in ONS cells derived from healthy control (HC), MCI and AD individuals. We identified AD- and MCI-associated proteomic alterations that may provide insight into disease-associated molecular pathways and highlight candidate biomarker signatures for further validation.

## 3. Results

### 3.1. ONS cells from AD, MCI and HC individuals show comparable immature neuroglial characteristics

OM cells derived from nasal biopsies of individuals with AD, MCI and HC were used to generate olfactory neurospheres and ONS cell cultures using our previously established protocol [23,27]. The derivation, growth characteristics and immature neuroglial phenotype of this patient-derived model, including disease-associated transcriptomic features, have been described previously [23]. For the present proteomics study, cultures were assessed before sample collection to confirm that they retained the expected morphology and neuroglial marker profile.

Neurospheres from all diagnostic groups showed the expected compact, rounded morphology, consistent with our previous observations (**Fig. 1B**). Following neurosphere formation, spheres were dissociated and expanded as adherent ONS cells in monolayer culture. These cells displayed the typical elongated, spindle-shaped morphology and grew as adherent cultures under serum-containing conditions (**Fig. 1C**).

To confirm the phenotype of the ONS cultures used for proteomics profiling, expression of stem cell-, neuronal-, and glial-associated markers was assessed by qRT-PCR and immunocytochemistry. qRT-PCR analysis showed expression of the stem/progenitor markers Nestin and PAX6, immature neuronal markers βIII-tubulin (*TUBB3*) and microtubule-associated protein 2 (*MAP2*), and the glial-associated marker S100B across HC, MCI and AD ONS cell lines (**Fig. 1D**). Immunocytochemistry further confirmed expression of the stem cell marker Nestin, neuronal markers TUBB3 and doublecortin (DCX), and astroglial-associated markers aquaporin 4 (AQP4) and S100B (**Fig. 1E**). Together, these morphological and molecular features indicate that the ONS cultures used in this study retained an immature neuroglial phenotype consistent with our previous characterisation of this patient-derived model.

### 3.2. Subcellular proteomics identifies compartment-associated protein abundance changes in AD and MCI ONS cells

#### 3.2.1. Overview of cytoplasmic and nuclear proteomic profiles

To improve proteome coverage and investigate whether disease-associated protein changes differed between cellular compartments, ONS cells derived from HC individuals and patients with MCI and AD were separated into nuclear and cytoplasmic fractions prior to proteomic analysis. Subcellular fractionation can increase the depth of proteomic profiling by reducing sample complexity and enriching proteins from distinct cellular compartments, while also providing information on compartment-associated protein abundance patterns [29,30]. This approach is also biologically relevant because altered nucleocytoplasmic transport and nuclear pore complex dysfunction have been reported in AD and tau-related neurodegenerative disease [31]. Although the present study did not directly assess nucleocytoplasmic transport, comparison of cytoplasmic and nuclear protein fractions may provide useful information on compartment-associated protein abundance patterns.

Data quality assessment showed broadly comparable protein intensity distributions before and after normalisation, supporting the suitability of four HC, six MCI and five AD ONS lines from both fractions for downstream analysis (**Fig. S1A, B**). Principal component analysis (PCA) of the cytoplasmic and nuclear fractions showed partial separation of AD ONS lines from HC and MCI samples, while HC and MCI samples showed greater overlap in both fractions (**Fig. 2A, B**).

**Fig. 2.**
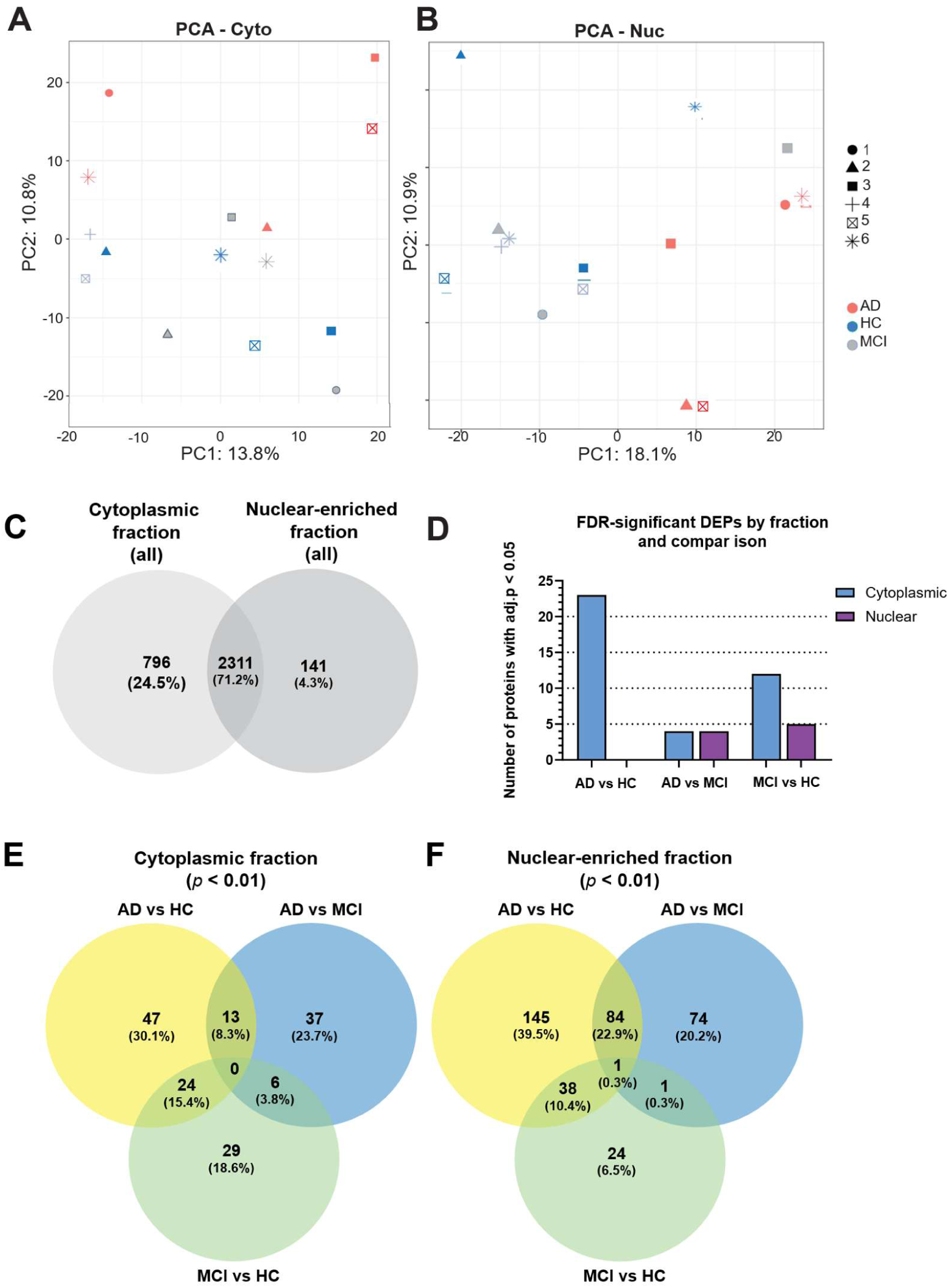
Overview of cytoplasmic and nuclear-enriched proteomic profiles in HC, MCI and AD ONS cells. (**A, B**) Principal component analysis (PCA) plots of (**A**) cytoplasmic and (**B**) nuclear-enriched proteins showing the distribution of HC, MCI and AD samples. AD samples are shown in red, HC samples in blue and MCI samples in grey. (**C**) Venn diagram showing the number and percentage of shared and unique proteins detected in the cytoplasmic and nuclear fractions. (**D**) Bar graph showing the number of FDR-significant differentially expressed proteins (DEPs) identified in each comparison within the cytoplasmic and nuclear fractions. FDR-significant proteins were defined using an adjusted *p*-value < 0.05. (**E, F**) Venn diagrams showing the overlap of exploratory candidate proteins identified using a nominal threshold of *p* < 0.01 across AD vs HC, AD vs MCI and MCI vs HC comparisons in the (**E**) cytoplasmic and (**F**) nuclear fractions. PCA plots and differential abundance analyses were generated using LFQ-Analyst.

In total, 3,248 proteins were identified across all samples in the cytoplasmic and nuclear fractions, including 3,107 proteins in the cytoplasmic fraction and 2,452 proteins in the nuclear fraction. Most proteins were detected in both fractions, while smaller subsets were detected only in the cytoplasmic and nuclear fractions (**Fig. 2C**).

Differential protein abundance analysis was conducted across three pairwise comparisons: AD vs HC, AD vs MCI, and MCI vs HC. Differentially abundant proteins (DEPs) were defined using FDR-adjusted *p*-values to identify the most robust protein changes after multiple-testing correction. Proteins meeting a nominal significance threshold of *p* < 0.01, but not remaining significant after FDR correction, were considered exploratory candidate proteins.

The largest number of significant DEPs was observed in the cytoplasmic AD vs HC comparison (**Fig. 2D**). In the nuclear-enriched fraction, no proteins remained significant after FDR correction in the AD vs HC comparison, whereas significant DEPs were detected in the AD vs MCI and MCI vs HC comparisons (**Fig. 2D**). Given the limited number of significant proteins in some comparisons, we next examined exploratory candidate proteins to assess broader disease- and compartment-associated proteomic patterns. At the nominal *p* < 0.01 level, the broadest changes were observed in the nuclear-enriched fraction, particularly in the AD vs HC and AD vs MCI comparisons (**Fig. 2E, F**). Therefore, throughout the following sections, FDR-significant DEPs are interpreted as the strongest individual findings, while nominally significant candidate proteins are considered exploratory candidates that may support pathway-level interpretation and require further validation.

#### 3.2.2. Selected disease-associated protein changes and western blot assessment

To further examine selected disease-associated protein abundance changes, representative comparisons were selected primarily based on the presence of proteins that remained significant after FDR correction, together with biological relevance and overlap with reconstructed proteomics findings. Volcano plots were generated for the nuclear-enriched AD vs MCI, cytoplasmic AD vs HC and cytoplasmic MCI vs HC comparisons (**Fig. 3A–C**). In these plots, proteins that remained significant after FDR correction were highlighted as the most robust findings, while proteins meeting the nominal *p* < 0.01 threshold were shown to visualise broader abundance patterns. Heatmaps were then used to display proteins that remained significant after FDR correction in the nuclear-enriched and cytoplasmic fractions (**Fig. 3D, E**). Full differential abundance results for all DEPs and selected exploratory candidate proteins described in this section are provided in Table S1. Volcano plots for the remaining fraction-specific comparisons are provided in Fig. S2.

**Fig. 3.**
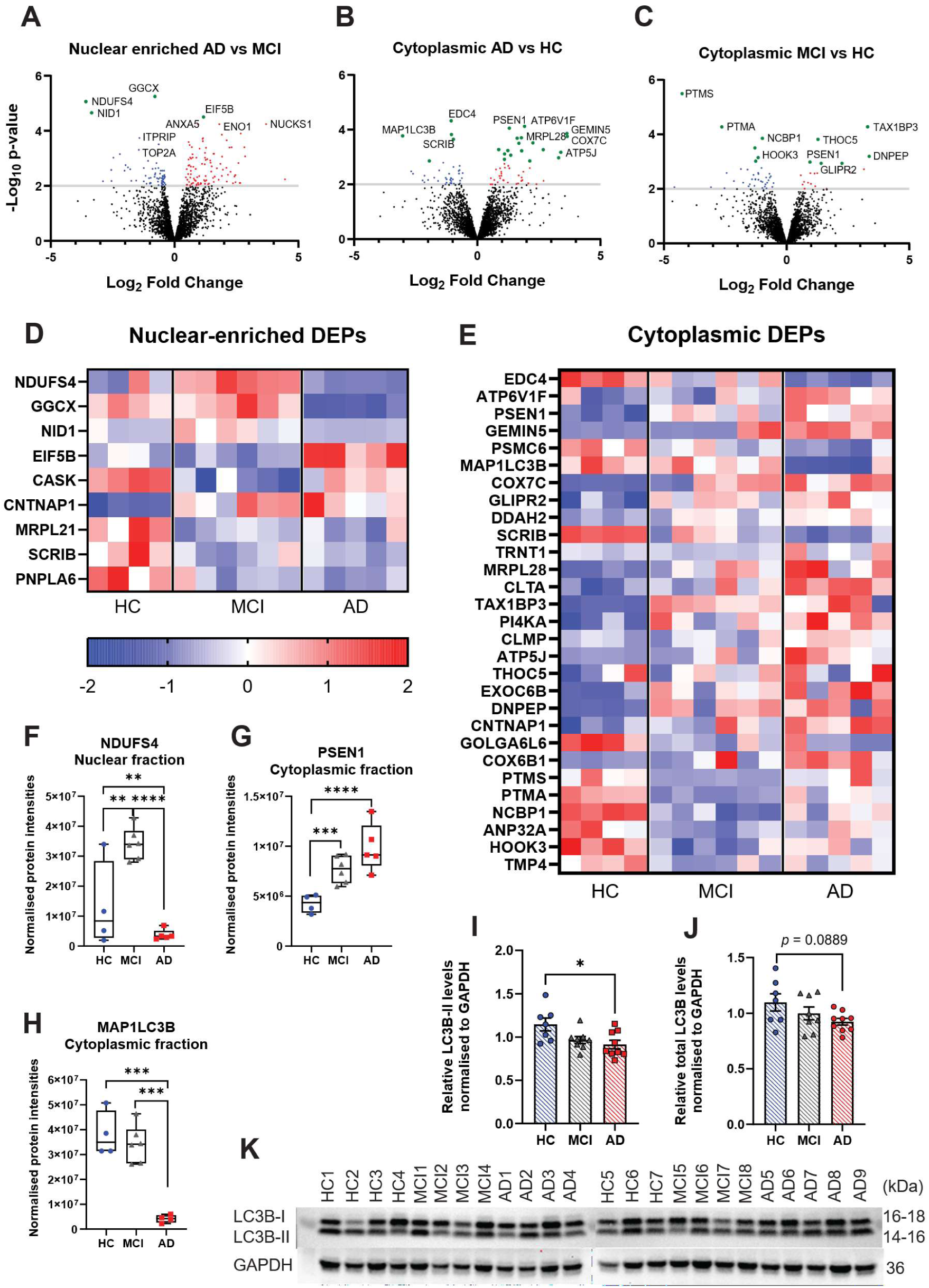
Selected cytoplasmic and nuclear-enriched proteomic alterations in AD and MCI ONS cells. Volcano plots showing differentially abundant proteins in (**A**) the nuclear-enriched AD vs MCI, (**B**) the cytoplasmic AD vs HC, and (**C**) the cytoplasmic MCI vs HC comparisons. Green points indicate proteins that remained significant after FDR correction, while red and blue points indicate proteins meeting the nominal significance threshold of *p* < 0.01 according to direction of change. Nominally significant proteins are shown in the volcano plots to visualise broader abundance patterns, whereas proteins significant after FDR correction are highlighted as the most robust findings. (**D, E**) Heatmaps showing row-scaled abundance of DEPs that remained significant after FDR correction in the nuclear-enriched and cytoplasmic fractions. Log_2_-transformed imputed protein intensity values were z-score scaled across samples within each fraction. Red indicates relatively higher abundance, and blue indicates relatively lower abundance for each protein. (**F-H**) Boxplots showing normalised protein intensities for selected representative proteins. (**I-K**) Western blot analysis of MAP1LC3B, commonly referred to as LC3B, in HC (*n* = 7), MCI (*n* = 8) and AD (*n* = 9) ONS cells across two independent western blot experiments. (**I**) LC3B-II was significantly reduced in AD ONS cells compared with HC. (**J**) Total LC3B showed a similar, non-significant trend. (**K**) Representative LC3B and GAPDH western blot image from one of the two independent experiments. Band intensities were normalised to GAPDH. Data are shown as mean ± SEM. \**p* < 0.05.

In the nuclear-enriched fraction, AD- and MCI-associated changes included proteins linked to mitochondrial and metabolic pathways, translation, ECM-associated biology, cell adhesion, cytoskeletal organisation, cell polarity and lipid metabolism. Among these, NADH: ubiquinone oxidoreductase subunit S4 (NDUFS4), a Complex I-associated protein, showed one of the clearest diagnostic-group-associated abundance patterns and remained significant after FDR correction in the AD vs MCI comparison and was also nominally significant in the AD vs HC and MCI vs HC comparisons. NDUFS4 was reduced in AD compared with both HC and MCI, while being increased in MCI compared with HC (**Fig. 3A, D, F**). Other NADH dehydrogenase subunits, including NDUFS7 and NDUFS6, showed nominally significant changes, suggesting that additional Complex I-associated proteins may contribute to the broader proteomic pattern. Exploratory changes were also observed in other mitochondrial- and metabolism-associated proteins, including Complex IV-associated subunits, ATP synthase- or proton transport-associated proteins, glycolysis-associated proteins, and proteins involved in mitochondrial transport or metabolism.

In addition to mitochondrial- and metabolism-associated changes, several proteins that remained significant after FDR correction were associated with other biological functions. GGCX, an enzyme involved in vitamin K-dependent protein carboxylation, a pathway that has been linked to brain health and cognitive function [32–34] was the most significant nuclear-enriched DEP in the AD vs MCI comparison after FDR correction and was consistently reduced in AD compared with both MCI and HC (**Fig. 3A, D and Fig. S2A**). In contrast, GGCX abundance was not significantly altered between cohorts in the cytoplasmic fraction, suggesting that this change was specific to the nuclear-enriched proteome. Additional FDR-significant proteins in the nuclear AD vs MCI comparison included NID1, an extracellular matrix and basement membrane-associated protein [35] and EIF5B, a translation initiation factor involved in ribosomal subunit joining [36] (**Fig. 3A, D**).

In the nuclear MCI vs HC comparison, CNTNAP1, a neurexin-family protein involved in neuronal adhesion [37] was significantly increased, whereas SCRIB, a scaffolding protein involved in cell polarity and cell adhesion [38] and PNPLA6, a phospholipase-associated protein involved in lipid metabolism [39], were significantly reduced in MCI compared with HC (**Fig. 3D and Fig. S2B**). CASK, a scaffolding protein associated with neuronal signalling and synaptic organisation [40] was also reduced in MCI compared with both HC and AD (**Fig. 3D and Fig. S2B**).

In the cytoplasmic fraction, several DEPs showed altered abundance in AD and/or MCI ONS cells compared with HC (**Fig. 3B, C, E and Fig. S2C**). PSEN1, a catalytic component of the γ-secretase complex involved in APP processing and familial AD [41], was significantly increased in both AD and MCI compared with HC and was among the most significant proteins differentiating disease groups from controls (**Fig. 3B, C, E, G**). EDC4, a scaffold protein involved in mRNA decapping and processing body biology [42], MAP1LC3B, an autophagy-associated protein [43] and SCRIB were significantly reduced in AD compared with HC and/or MCI (**Fig. 3B, E, H and Fig. S2C**). PTMS and PTMA, proteins linked to chromatin organisation and transcriptional regulation [44], were also among the most significant cytoplasmic DEPs showing diagnostic-group-associated abundance differences, with the strongest reductions observed in MCI compared with HC and higher protein levels in AD compared with MCI (**Fig. 3C, E**). HOOK3, a protein involved in intracellular trafficking and microtubule-associated transport [45], showed a similar pattern (**Fig. 3C, E**).

Because MAP1LC3B, commonly referred to as LC3B, was one of the most significantly reduced cytoplasmic proteins in AD ONS cells, LC3B was further assessed by western blot. LC3B-II/GAPDH was significantly reduced in AD ONS cells compared with HC, whereas total LC3B/GAPDH showed a similar but non-significant trend (**Fig. 3I–K**). These findings provide supportive evidence for reduced LC3B abundance in AD ONS cells but do not establish altered autophagic flux. LC3B-I/LC3B-II and LC3B-II/LC3B-I ratios were also examined as additional autophagy-associated measures, however, no consistent group-level differences were observed across HC, MCI and AD ONS cells (**Fig. S3A, B**). Western blot analysis was also attempted for additional selected candidates, including NDUFS4 and PSEN1, but these experiments did not provide significant or consistent support for the proteomics findings and were therefore not included as main validation panels. Immunofluorescence analysis was also performed for selected candidates, including PSEN1, ENO1, EDC4 and LC3B, but did not reveal clear differences in nuclear or cytoplasmic localisation between groups. Full LC3B membranes and corresponding GAPDH membranes are shown for experiment 1 (**Fig. S3C, D**) and experiment 2 (**Fig. S3E, F**).

### 3.3. Fraction-specific proteomic enrichment analysis identifies metabolic, cytoskeletal, transcriptional and RNA-processing alterations in AD and MCI ONS cells

To explore the biological pathways associated with differentially expressed proteins, PANTHER enrichment analysis was performed using proteins altered at *p* < 0.05 in each comparison. This analysis focused on PANTHER GO-slim biological processes and protein classes. Across the cytoplasmic fraction, 88 biological processes and 20 protein classes were significantly enriched (FDR < 0.05). Corresponding analysis of the nuclear fraction identified 158 significantly enriched biological processes and 33 significantly enriched protein classes.

In the cytoplasmic fraction, enriched protein classes and biological processes were mainly associated with transcriptional regulation, oxidoreductase activity, cytoskeletal organisation, extracellular matrix (ECM) proteins and cellular metabolic or catabolic processes (**Fig. 4A–C**). In AD vs HC cytoplasmic proteins, the most enriched categories included DNA-binding transcription factors, oxidoreductases, actin filament organisation and cellular macromolecule catabolic processes (**Fig. 4A**). The AD vs MCI comparison showed enrichment of DNA-binding transcription factors, metabolic interconversion enzymes, chaperones, oxidoreductases, ECM proteins and cytoskeletal proteins (**Fig. 4B**). In the MCI vs HC comparison, enriched categories included RNA splicing and RNA processing factors, DNA-binding transcription factors, RNA metabolism proteins and actin-binding/cytoskeletal proteins (**Fig. 4C**).

**Fig. 4.**
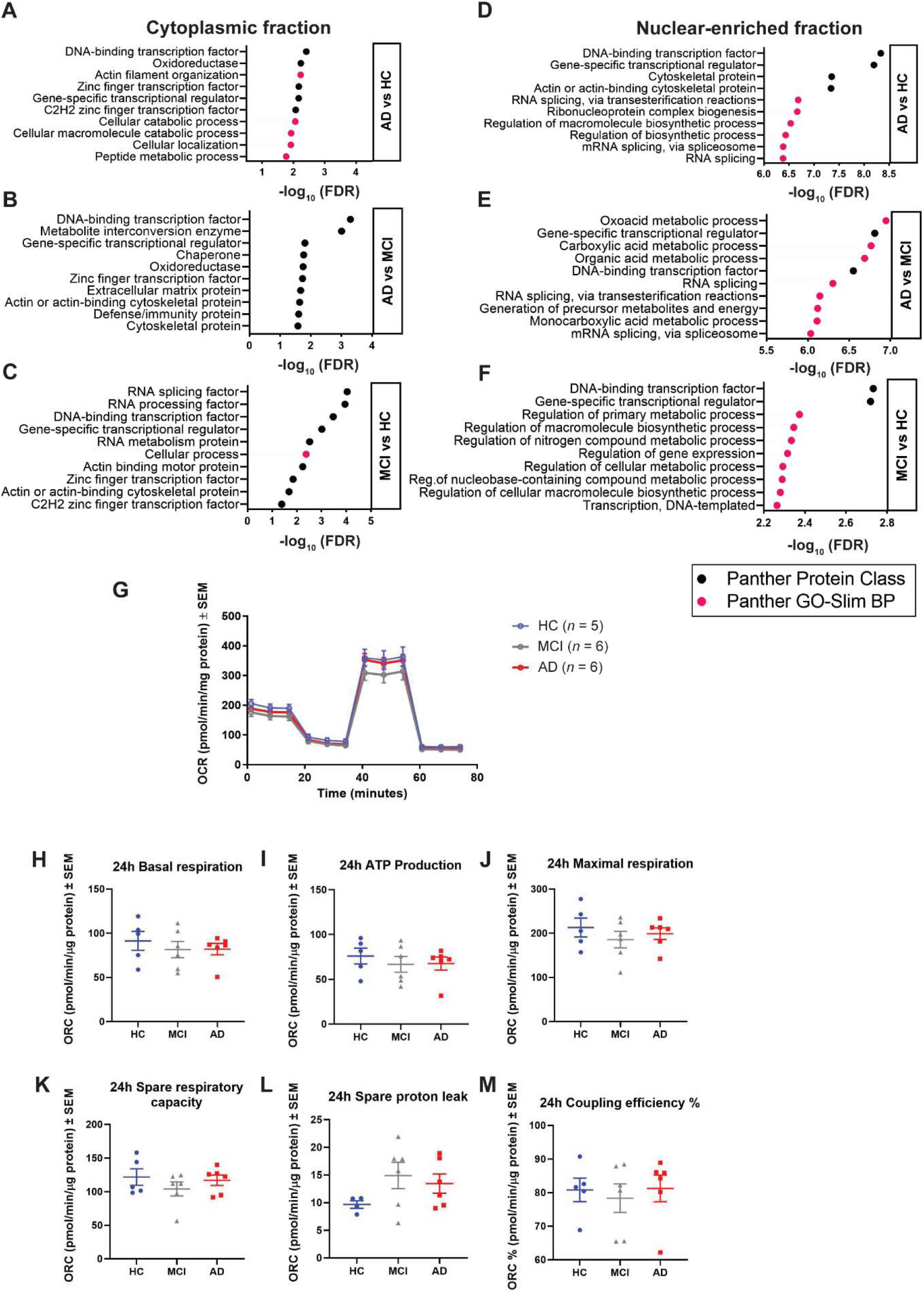
Fraction-specific proteomic enrichment analysis and assessment of mitochondrial respiration in HC, MCI and AD ONS cells. **(A-F)** PANTHER enrichment analysis of proteins altered at *p* < 0.05 in the cytoplasmic and nuclear-enriched fractions. The most significantly enriched PANTHER protein classes and PANTHER GO-Slim biological processes (Panther GO-Slim BP) are shown in the cytoplasmic fraction in (**A**) AD vs HC, (**B**) AD vs MCI and (**C**) MCI vs HC comparisons, and for the nuclear-enriched fraction in (**D**) AD vs HC, (**E**) AD vs MCI and (**F**) MCI vs HC comparisons. Black dots indicate enriched PANTHER protein classes and pink dots indicate enriched PANTHER GO-Slim biological processes. Enrichment significance is shown as -log10(FDR), with higher values indicating stronger enrichment. (**G-M**) Mitochondrial respiration was assessed using the Seahorse XF Cell Mito Stress Test in HC (*n* = 5), MCI (*n* = 6) and AD (*n* = 6) ONS cell lines. **(G)** Representative oxygen consumption rate (OCR) profile showing OCR measurements at baseline and following sequential injections of oligomycin, FCCP, and rotenone/antimycin A. OCR values are expressed as pmol/min/μg protein. **(H)** Basal respiration, **(I)** ATP production, **(J)** Maximal respiration, **(K)** Spare respiratory capacity, **(L)** Proton leak and **(M)** Coupling efficiency. Each symbol in H-M represents an individual donor, calculated from five technical replicate wells. Data are presented as mean ± SEM. One HC donor value was excluded from the proton leak analysis following Grubbs’ test (α = 0.05); no other biological data points were excluded. Statistical comparisons among groups were performed using one-way ANOVA followed by Tukey’s multiple comparisons test. No significant group-level differences were detected for any respiratory parameter.

In the nuclear fraction, enrichment analysis revealed a stronger association with transcriptional regulation, RNA splicing, ribonucleoprotein complex biogenesis, metabolic processes and regulation of biosynthetic processes (**Fig. 4D–F**). In the AD vs HC comparison, enriched terms included DNA-binding transcription factors, gene-specific transcriptional regulators, cytoskeletal proteins, RNA splicing and ribonucleoprotein complex biogenesis (**Fig. 4D**). The AD vs MCI nuclear comparison showed enrichment for oxoacid, carboxylic acid and organic acid metabolic processes, DNA-binding transcription factors, RNA splicing and generation of precursor metabolites and energy (**Fig. 4E**). In the MCI vs HC comparison, enriched nuclear categories included DNA-binding transcription factors, gene-specific transcriptional regulators, regulation of primary and macromolecule biosynthetic processes, nitrogen compound metabolism and cellular metabolic processes (**Fig. 4F**).

### 3.4. Assessment of mitochondrial respiration in AD ONS cells

Given the enrichment of metabolic and mitochondrial-associated proteins and pathways identified in the proteomic analyses, we next investigated whether these molecular alterations were associated with changes in mitochondrial function. Cellular respiration was assessed in HC, MCI and AD ONS cells using the Seahorse XF Cell Mito Stress Test. Overall oxygen consumption rate (OCR) profiles were similar across the three groups (**Fig. 4G**). Quantification of key respiratory parameters revealed no significant differences among the groups in basal respiration, ATP-linked respiration, maximal respiration, spare respiratory capacity, proton leak, or coupling efficiency (**Fig. 4H-M**). Although minor group-level trends were observed, including slightly lower ATP-linked respiration in MCI and AD (**Fig. 4I**) and modestly increased proton leak in MCI and AD (**Fig. 4L**), these differences were variable and did not reach statistical significance.

### 3.5. Reconstructed bulk proteomics identifies an AD-associated protein signature enriched for extracellular matrix, cytoskeletal and organelle-associated pathways

Next, we performed reconstructed bulk proteomics analysis to provide an integrated view of proteomic changes across the nuclear-enriched and cytoplasmic fractions of the same ONS cell samples analysed in the fractionated proteomics workflow (**Fig. S4**). A total of 2,490 proteins were detected in the reconstructed bulk dataset. PCA of bulk proteomic profiles showed partial group-level separation, with AD samples clustering apart from healthy controls along PC1, which accounted for 37.18% of the variance (**Fig. 5A**). MCI samples showed a more variable distribution and overlapped with both AD and HC groups, whereas HC samples clustered more closely together, suggesting lower within-group variation.

**Fig. 5.**
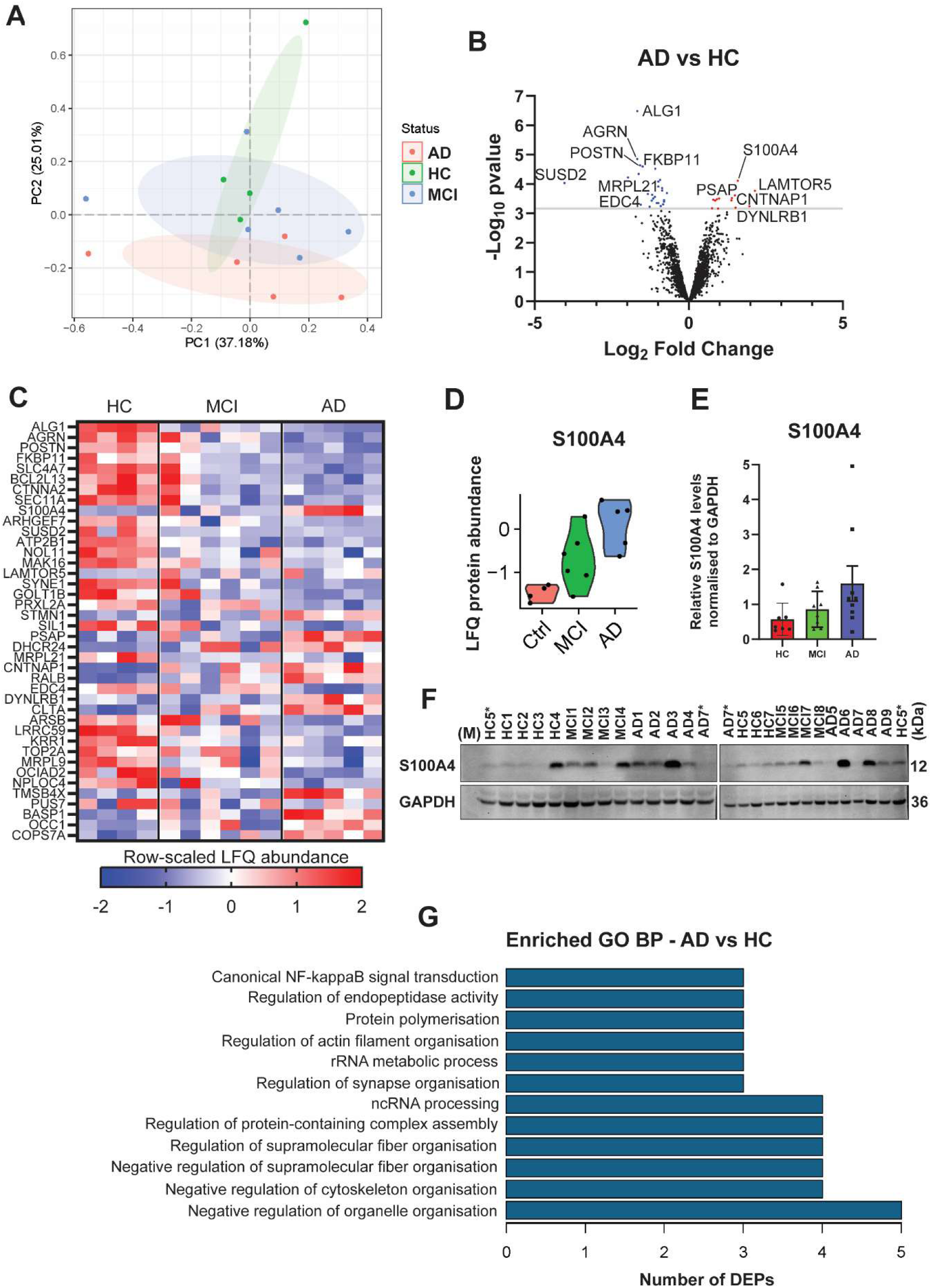
Bulk proteomics identifies an AD-associated protein signature in ONS cells. **(A)** PCA plot showing the distribution of HC, MCI and AD ONS samples based on reconstructed bulk LFQ proteomics profiles. **(B)** Volcano plot showing differentially abundant proteins in AD compared with HC. Proteins passing FDR < 0.05 are highlighted and represented in red (upregulated in AD) and blue (downregulated in AD). **(C)** Heatmap showing row-scaled LFQ abundance values for the 40 FDR-significant AD vs HC DEPs across HC, MCI and AD samples. **(D)** Violin plot showing S100A4 LFQ protein abundance across HC, MCI and AD samples. **(E)** Western blot quantification of S100A4 in HC (n = 7), MCI (n = 8) and AD (n = 9) ONS cells across three independent western blot experiments, normalised to GAPDH and expressed relative to the average S100A4/GAPDH value across all biological samples. Data are shown as mean ± SEM. **(F)** Representative S100A4 and GAPDH western blot image from experiment 2. HC5* and AD7* were included on both membranes as bridging samples. **(G)** Selected enriched GO biological process terms from the 40 FDR-significant AD vs HC reconstructed bulk proteomics DEPs, identified using ClusterProfiler. The x-axis indicates the number of DEPs contributing to each enriched term. Differential abundance was assessed using limma moderated t-tests with Benjamini–Hochberg correction.

Differential protein abundance analysis identified 40 proteins in AD ONS cells compared with HC after FDR correction, using an adjusted *p* < 0.05 threshold. Of these, 27 proteins were decreased, and 13 proteins were increased in AD compared with HC (**Fig. 5B, C and Table S2**). Compared with the fraction-specific analyses, the reconstructed bulk dataset identified a smaller but more integrated AD-associated protein signature, likely reflecting protein changes that remained evident when cytoplasmic and nuclear-enriched signals were combined. The most significantly altered proteins included ALG1, which is involved in protein glycosylation [46] AGRN and POSTN, extracellular matrix-associated proteins [47,48], FKBP11 and SEC11A, proteins linked to ER/protein processing [49,50], SLC4A7, an ion transporter [51], BCL2L13, a mitochondrial-associated protein [52], CTNNA2, a cytoskeletal and cell adhesion-associated protein [53] and S100A4, a calcium-binding protein linked to cell motility, stress responses and inflammatory signalling [54].

Only one protein, DYNLRB1, was significantly altered between AD and MCI samples, and no proteins were significantly altered between MCI and HC samples (**Fig. S5A, B**). DYNLRB1 is a component of the cytoplasmic dynein motor complex involved in microtubule-based intracellular transport [55]. DYNLRB1 was also significantly increased in AD compared with HC, suggesting that this protein may contribute to the AD-associated reconstructed bulk proteomic signature.

Several proteins identified in the reconstructed bulk dataset also overlapped with the fraction-specific analyses. EDC4, CNTNAP1 and CLTA overlapped with FDR-significant cytoplasmic DEPs. Additional overlapping proteins included BASP1, TMSB4X, BCL2L13, POSTN, PSAP, SEC11A, NPLOC4 and DYNLRB1, which were detected as nominal candidates in both cytoplasmic and nuclear-enriched fractions. Conversely, several proteins that were nominally altered in the reconstructed bulk dataset reached FDR significance in the fraction-specific analyses, including PTMS, PNPLA6, PSEN1 and SCRIB.

S100A4 was selected for western blot analysis based on its biological relevance and significant increase in the reconstructed bulk proteomics dataset (**Fig. 5B-D**). Across three independent western blot runs, S100A4 showed a reproducible trend towards increased abundance in AD ONS cells, consistent with the direction of change observed by proteomics (**Fig. 5E, F**). However, this finding did not reach statistical significance, and individual variability was observed between patient-derived ONS cell lines. Full western blot membranes for S100A4 are shown in Fig. S6.

GO enrichment analysis of the 40 significantly altered AD vs HC bulk proteomics DEPs further supported changes in ECM, cytoskeletal, ribosomal/mitochondrial and lysosomal-associated pathways. Selected enriched biological process terms included regulation of cytoskeletal and supramolecular fibre organisation, regulation of organelle organisation, RNA/rRNA processing, protein polymerisation, synapse organisation, endopeptidase activity and canonical NF-κB signal transduction (**Fig. 5G**). The full GO biological process enrichment plot, together with additional cellular component and molecular function enrichment terms, is provided in Fig. S7.

### 3.6. Integrated transcriptomic and proteomic analysis identifies AD-associated RNA-protein correlation networks

To further assess concordance between the reconstructed bulk proteomics findings and transcriptomic changes previously identified in the same ONS cohort [23], we performed RNA-protein correlation analysis on genes detected across both datasets. The reconstructed bulk proteomics dataset contained 2,490 proteins in total. Of the 16,585 protein-coding transcripts detected in the RNA-seq dataset, 1,249 had corresponding protein-level data in the reconstructed bulk proteomics dataset, reflecting partial coverage of the transcriptome by the proteomics dataset (**Fig. 6A**). For genes with multiple isoforms or transcripts, the transcript with the highest median expression was retained. Correlation analysis identified 45 genes with significant transcript-protein relationships at *p* < 0.01, of which 41 showed positive correlations and 4 showed negative correlations (**Table S3**). These significant RNA-protein correlations represented only a subset of the matched dataset, indicating partial RNA-protein concordance.

**Fig. 6.**
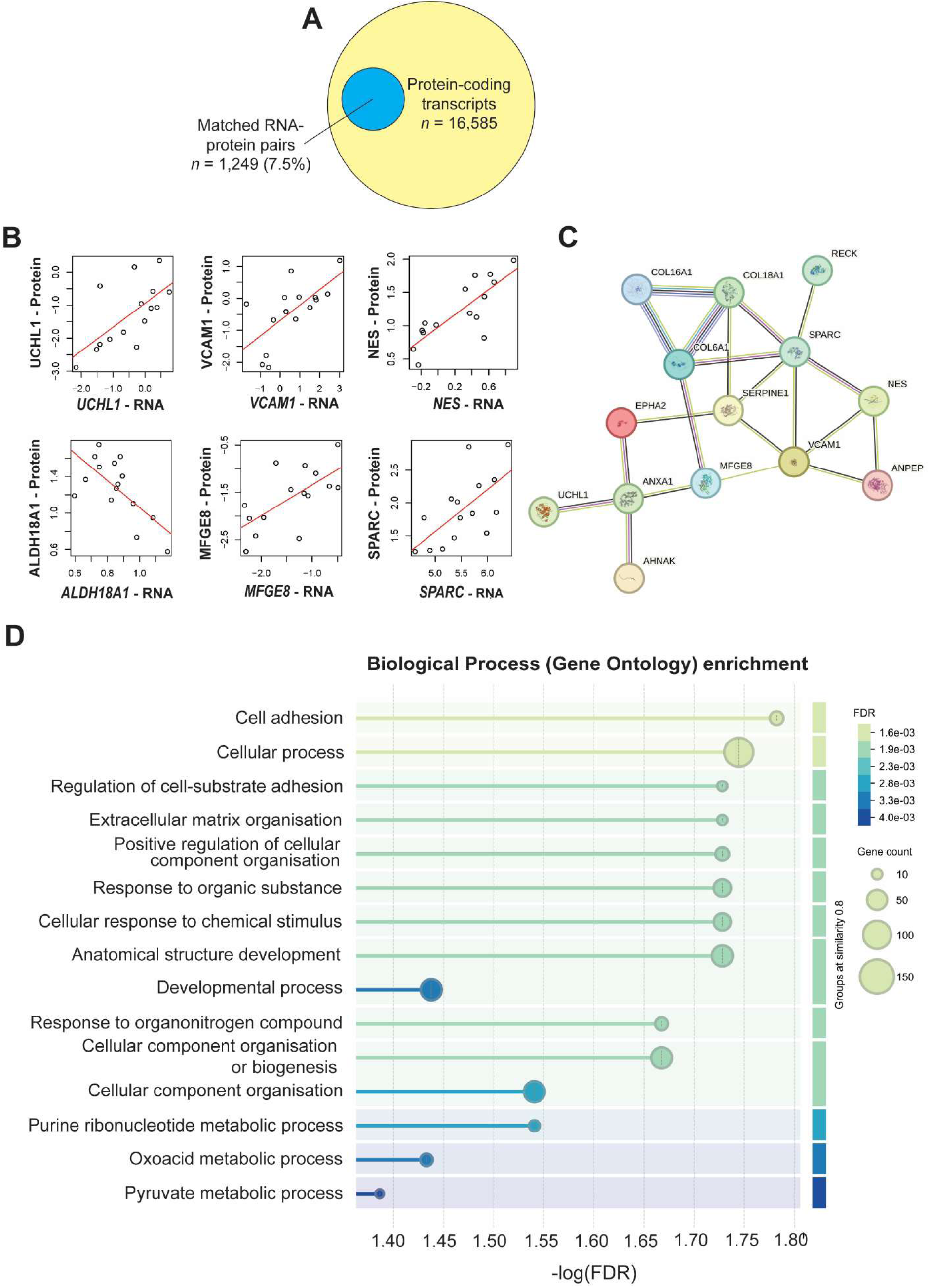
Integrated transcriptomic and proteomic analysis identifies AD-associated RNA–protein correlation networks. (**A**) Schematic showing overlap between transcriptomic and reconstructed bulk proteomic datasets. Of the 16,585 protein-coding transcripts detected at the RNA level, 1,249 had corresponding protein-level data in the reconstructed bulk proteomics dataset, representing 7.5% coverage of the protein-coding transcriptome by the proteomics dataset. (**B**) Representative RNA-protein correlation plots (*p* < 0.01) for selected genes detected in both datasets. (**C**) Protein–protein interaction network of significantly correlated RNA-protein pairs (*p* < 0.01) generated using the STRING database, showing functional connectivity among AD-associated proteins. (**D**) Gene Ontology biological process enrichment analysis of correlated genes at *p* < 0.05, showing the top 15 biological processes across HC, MCI and AD groups. Bubble size indicates gene count, and colour indicates FDR-adjusted enrichment significance (FDR < 0.05).

Several significantly correlated RNA-protein pairs represented functional processes relevant to AD and neurodegeneration. These included genes involved in ECM organisation and cell adhesion, such as *COL6A1*, *COL16A1*, *COL18A1*, *SPARC*, *VCAM1* and *RECK* [56–59], inflammatory or stress-related pathways, including *ANXA1*, *SERPINE1* and *MFGE8* [60–62], mitochondrial or metabolic regulation, including *NDUFV1* and *ALDH18A1* [63,64], and neuronal, cytoskeletal or proteostasis-related processes, including *UCHL1*, *NES*, *SDCBP* and *PFN2* [65–68]. Representative scatter plots demonstrated concordant transcript-protein relationships for selected genes, including *UCHL1*, *VCAM1*, *NES*, *MFGE8* and *SPARC*, whereas *ALDH18A1* showed a negative transcript-protein correlation (**Fig. 6B**).

To determine whether these correlated genes were functionally connected, we performed STRING interaction analysis using genes with transcript-protein correlations at *p* < 0.01. Interestingly, this analysis revealed an interconnected network involving AD-associated, ECM, adhesion, inflammatory/vascular and cytoskeletal-associated proteins, including COL18A1, COL16A1, COL6A1, SPARC, SERPINE1, VCAM1, MFGE8, ANXA1, EPHA2, UCHL1, RECK and NES (**Fig. 6C**). GO biological process enrichment analysis of correlated genes at *p* < 0.05 further supported this pattern, identifying enrichment for cell adhesion, ECM organisation, regulation of cell–substrate adhesion, cellular component organisation, response to chemical or organic stimuli, developmental processes, purine ribonucleotide metabolism, oxoacid metabolism and pyruvate metabolism (**Fig. 6D**). Additional functional enrichment analyses, including KEGG pathway, GO molecular function and GO cellular component analyses showed overlapping enrichment of ECM/adhesion-related and metabolic processes (**Fig. S8**). Together, these findings suggest that selected RNA-protein relationships align with the reconstructed bulk and fraction-specific proteomics findings at the pathway level, while the relatively small number of significant correlations indicates limited overall RNA-protein concordance across the matched dataset.

### 3.7. Summary of main molecular findings

A schematic summary of the main disease-associated molecular findings identified across the fraction-specific proteomics, reconstructed bulk proteomics and integrated transcriptomic– proteomic analyses is shown in Fig. 7.

**Fig. 7.**
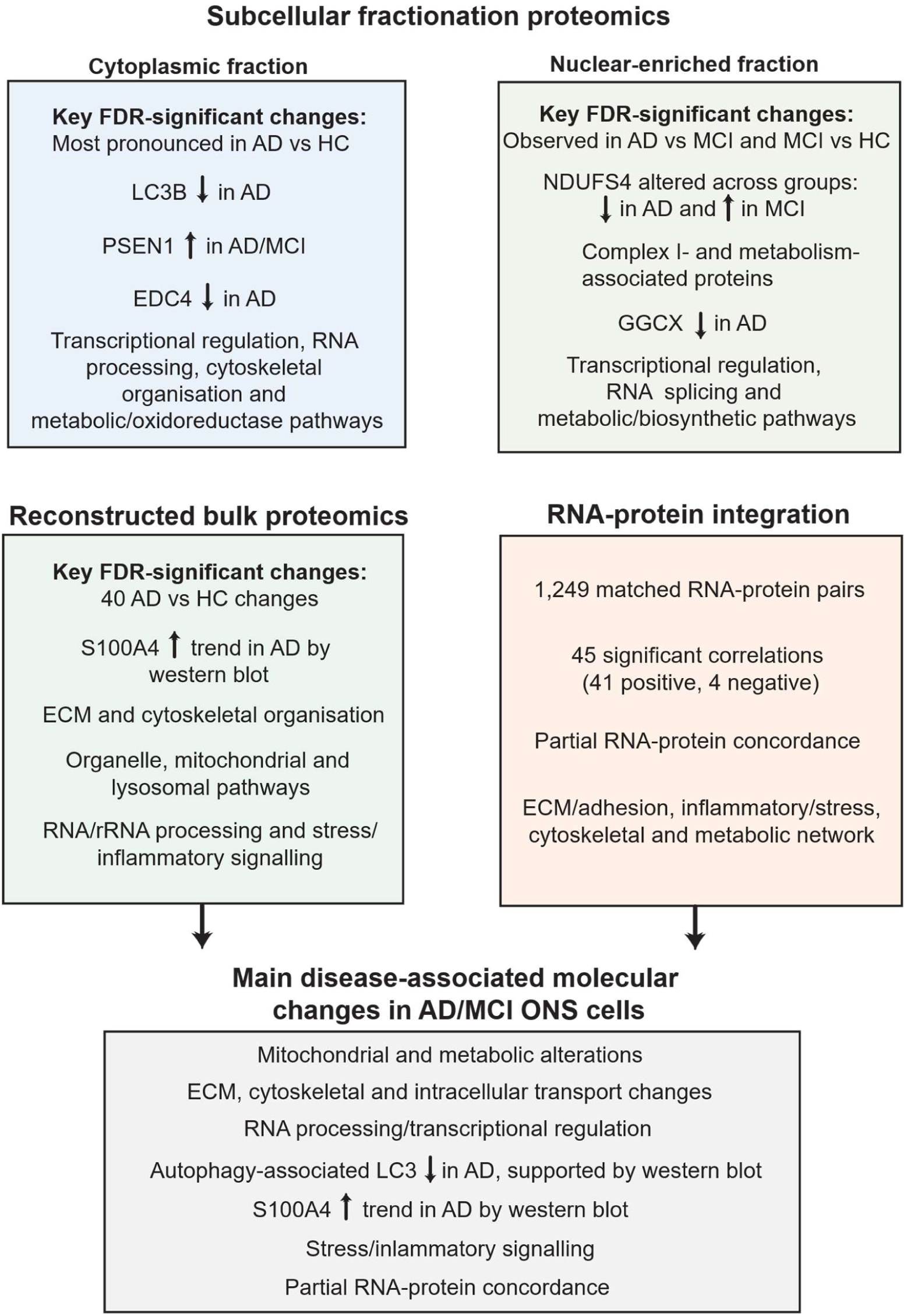
Summary of the main disease-associated molecular findings identified in AD and MCI ONS cells. Fraction-specific proteomics identified distinct cytoplasmic and nuclear-enriched protein abundance changes, including alterations in autophagy-associated, mitochondrial/metabolic, cytoskeletal, RNA-processing and transcriptional pathways. Reduced LC3B abundance in AD ONS cells was supported by western blot analysis.

Reconstructed bulk proteomics identified 40 FDR-significant proteins in AD compared with HC and highlighted ECM, cytoskeletal, organelle, mitochondrial, lysosomal, RNA/rRNA-processing and stress/inflammatory pathways, with S100A4 showing a reproducible trend toward increased abundance by western blot. Integrated RNA–protein analysis identified 1,249 matched RNA–protein pairs and 45 significant correlations, indicating partial RNA–protein concordance and functional connectivity among ECM/adhesion, inflammatory/stress, cytoskeletal and metabolic processes.

## 4. Discussion

In the present study, subcellular fractionation followed by bottom-up proteomics was used to investigate protein abundance changes in patient-derived ONS cells from HC, MCI and AD individuals. A key strength of this approach is the use of patient-derived cells, which retain donor-specific biological variation and provide a clinically relevant model for examining molecular changes associated with AD and MCI. At the same time, this patient-derived nature introduces heterogeneity between cell lines, which may contribute to variability across individual samples. Rather than reflecting only technical variation, this heterogeneity may also represent real biological differences between individuals. Therefore, larger cohorts will be required to distinguish robust disease-associated signatures from donor-specific variation.

The ONS cultures used in this study showed the expected neurosphere-forming capacity, adherent spindle-shaped morphology and expression of stem cell-, neuronal- and glial-associated markers, consistent with our previous characterisation of patient-derived ONS cells [23]. These findings support the use of ONS cells as an accessible, patient-derived *in vitro* model for investigating molecular changes associated with AD and MCI. However, as ONS cells are immature neuroglial-like cells rather than mature neurons or glia, these findings should be interpreted as disease-associated changes in an olfactory stem cell-derived model rather than direct representations of mature brain cell pathology.

The subcellular fractionation approach provided complementary cytoplasmic and nuclear-enriched proteomic profiles. However, substantial overlap between fractions was observed, which was expected, as the cellular proteome is dynamic, with many proteins continuously moving between the cytoplasm and the nucleus [69]. In addition, because membrane-associated and organelle-derived proteins were not fractionated separately, the cytoplasmic fraction was expected to contain proteins from multiple cellular compartments [70]. Marker protein analysis supported enrichment of the nuclear and cytoplasmic fractions, however, mitochondrial resident proteins were detected in both fractions (**Fig. S9**). Therefore, fraction-specific findings should be interpreted as enrichment-based rather than representing completely isolated cellular compartments.

Differential protein abundance analysis identified clear disease-associated changes in ONS cells. When FDR-adjusted *p*-values were used to identify the most robust protein changes, the largest number of significant DEPs was observed in the cytoplasmic AD vs HC comparison (**Fig. 2D**), highlighting a clear AD-associated proteomic shift in ONS cells. Significant DEPs were also detected in the nuclear-enriched fraction in the AD vs MCI and MCI vs HC comparisons, suggesting possible disease-stage-associated differences in the nuclear-enriched proteome. When a broader nominal threshold of *p* < 0.01 was applied, the nuclear-enriched fraction contained a larger number of altered proteins than the cytoplasmic fraction, indicating that this fraction may capture broader disease-associated changes that do not all remain significant after correction for multiple testing.

In both cytoplasmic and nuclear fractions, the largest number of nominally significant candidate DEPs was identified in AD ONS cells compared with HC. This is consistent with the expectation that AD cells would show more pronounced proteomic alterations than healthy control cells [71]. The second highest number of candidate DEPs was observed in the AD vs MCI comparison, suggesting that proteomic differences were also evident between these diagnostic groups. However, as these cells were derived from different individuals rather than longitudinal samples from the same donors, these differences should not be interpreted as direct evidence of disease progression. Fewer candidate DEPs were observed between MCI and HC, indicating that the MCI proteomic phenotype may be more subtle or heterogeneous in this ONS cell cohort. The limited overlap of candidate DEPs across AD, MCI and HC comparisons further suggests that AD- and MCI-associated proteomic changes were largely distinct, although this pattern could also reflect cohort and patient heterogeneity. This is consistent with our previous RNA-seq findings in ONS cells, where fewer than 5% of differentially expressed genes were detected concomitantly when comparing AD and MCI with HC [23].

A notable feature of the fraction-specific dataset was the altered abundance of metabolic and mitochondrial-associated proteins. These included proteins linked to respiratory chain complexes, ATP synthase- or proton transport-associated processes, glycolysis, mitochondrial translation, calcium handling, protein import and intermediary metabolism (**Table S1**). NDUFS4 was one of the most notable candidates, showing diagnostic-group-associated differences in abundance together with changes in other NADH dehydrogenase subunits. NDUFS4 is a nuclear-encoded Complex I accessory subunit involved in Complex I assembly and stability [72,73] and altered Complex I activity has been reported in neurodegenerative disease contexts, including AD and related dementia brain tissue [74–76]. However, the direction and fraction-specific distribution of these changes varied between proteins, suggesting alterations in mitochondrial- and metabolism-associated protein abundance rather than a uniform reduction or activation of mitochondrial energy metabolism.

The detection of several mitochondrial-and metabolism-associated proteins in the nuclear-enriched fraction should be interpreted cautiously. Because the fractionation procedure enriches rather than isolates subcellular compartments, detection of these proteins in the nuclear-enriched fraction cannot establish definitive nuclear localisation. These findings may reflect altered subcellular distribution, stress-associated redistribution of mitochondrial proteins during mitochondrial import stress [77] or co-fractionation of organelle-associated material, including mitochondria, endoplasmic reticulum-associated proteins or mitochondria-associated membranes, with the nuclear-enriched fraction [78]. Therefore, although these changes may indicate broader compartment-associated proteomic differences in AD and MCI ONS cells, direct localisation studies will be required to determine their subcellular distribution.

To determine whether the proteomic changes were associated with altered mitochondrial respiratory function, mitochondrial respiration was assessed using the Seahorse Mito Stress Test. No significant differences were detected between HC, MCI and AD ONS cells under the basal assay conditions tested (**Fig. 4G-M**). This suggests that the observed mitochondrial- and metabolism-associated protein changes do not reflect overt respiratory dysfunction under these conditions. Instead, they may represent changes in protein regulation, subcellular localisation or other adaptive responses that occur without detectable changes in OCR-based respiration [79]. It is also possible that these changes occur upstream of measurable respiratory dysfunction, affect metabolic pathways not directly captured by OCR measurements, or become apparent only under additional metabolic stress, altered substrate availability, longer culture conditions or assays targeting specific respiratory chain complex activities.

Beyond mitochondrial and metabolic pathways, fraction-specific proteomics analysis identified selected proteins associated with autophagy, intracellular trafficking, RNA/P-body biology, cytoskeletal organisation, adhesion, ECM-associated processes and cellular stress responses. MAP1LC3B, commonly referred to as LC3B, was reduced in the cytoplasmic fraction of AD ONS cells, and western blot analysis showed a significant reduction in LC3B-II/GAPDH, while total LC3B/GAPDH showed a similar but non-significant trend. Because LC3B abundance alone does not reflect autophagic flux [80] these findings indicate altered abundance of an autophagy-associated protein rather than direct evidence of impaired autophagy. This is nevertheless of interest given previous reports implicating impaired autophagy and mitophagy in AD [81–83]. Other altered cytoplasmic and nuclear-enriched proteins, including PSEN1, EDC4, SCRIB, HOOK3, DYNLRB1, CASK, FRMD6, CNTNAP1, GGCX, NID1, EIF5B, PPIA, PTMS, PTMA and YWHAZ, further indicate that the proteomic differences identified in ONS cells extend beyond metabolic pathways to involve structural organisation, cellular regulation and intracellular transport.

Pathway enrichment analysis supported that the fraction-specific proteomic changes were organised around broader cellular networks rather than isolated individual proteins. In particular, enrichment patterns in the nuclear-enriched fraction suggested involvement of transcriptional and RNA-processing-related regulation, while cytoplasmic changes supported broader alterations in cellular structure, protein homeostasis and stress-associated processes. These findings are broadly consistent with our previous transcriptomic analysis of the same ONS cell cohort, which identified altered alternative splicing, cytoskeleton organisation, ECM organisation and cell adhesion pathways in AD and MCI ONS cells [23]. This comparison should be interpreted as pathway-level consistency between separate transcriptomic and proteomic analyses, rather than direct gene-protein concordance.

Reconstructed bulk proteomics analysis provided an integrated view of disease-associated protein changes across the combined cytoplasmic and nuclear datasets. In contrast to the fraction-specific analyses, which highlighted compartment-dependent changes, the reconstructed bulk analysis identified an overall AD-associated proteomic signature. Consistent with the fraction-specific results, the clearest changes were again observed in the AD vs HC comparison. PCA showed partial separation of AD samples from HC samples, while MCI samples showed a more variable and intermediate distribution. Differential abundance analysis identified 40 proteins significantly altered in AD compared with HC after FDR correction, while only one protein was significantly altered between AD and MCI and no proteins were significantly altered between MCI and HC. Together, these findings suggest that AD ONS cells display more evident proteomic alterations than HC cells, whereas MCI ONS cells may show a more subtle or heterogeneous proteomic phenotype. This is consistent with the clinical heterogeneity of MCI, as not all individuals with MCI progress to AD [84]. The lack of significant MCI vs HC findings may therefore reflect smaller effect sizes, greater biological variability within the MCI group, or limited statistical power due to the small cohort size.

GO enrichment analysis indicated that the AD-associated reconstructed bulk protein signature was enriched for structural, organelle-related and stress/inflammatory pathways. This pathway-level pattern was consistent with the fraction-specific proteomics results. DYNLRB1 was altered both in reconstructed bulk and fraction-specific datasets, supporting overlap between the analytical approaches and suggesting possible involvement of microtubule-based intracellular transport [85]. S100A4 was also significantly increased in the reconstructed bulk proteomics dataset and showed a reproducible, but non-significant, increase by western blot. S100A4 has been linked to AD-associated glial stress and inflammatory pathways, including altered astrocyte-mediated processing of neuron-derived mitochondria in aged 5xFAD mouse brain [86] and oxidative stress responses [87]. Although the proteomics finding identifies S100A4 as a candidate of interest, the western blot analysis did not provide statistically significant validation, and further studies will be required to determine its relevance in AD ONS cells.

Finally, the integrated RNA-protein analysis provided a more direct assessment of transcriptomic– proteomic alignment in the same ONS cell cohort. Whereas the earlier comparison between the current proteomic enrichment results and our previous RNA-seq findings suggested pathway-level overlap, direct RNA-protein correlation analysis identified a smaller subset of genes with coordinated transcript and protein abundance. This partial RNA-protein concordance is consistent with previous AD studies showing that disease-associated protein changes are not always directly reflected at the transcript level [88]. In AD, transcript and protein abundance may diverge due to post-transcriptional regulation, translational efficiency, protein stability, degradation and subcellular localisation, as well as technical differences between transcriptomic and proteomic platforms [89–91].

Despite this limited overlap, most significant RNA-protein relationships showed positive correlations, suggesting that a subset of disease-associated molecular changes in ONS cells may be transcriptionally coordinated. Many of these significantly correlated genes were linked to pathways also identified in the reconstructed bulk, fraction-specific and previous transcriptomic analyses, while the STRING network further highlighted functional connectivity among these processes. Together, these findings suggest that although direct RNA-protein concordance was limited, a subset of disease-associated molecular changes in ONS cells showed coordinated transcript-protein relationships within pathways relevant to AD-associated cellular dysfunction.

Overall, this study supports the presence of disease-associated proteomic alterations in patient-derived AD and MCI ONS cells. Subcellular fractionation highlighted compartment-associated changes, reconstructed bulk proteomics identified a clearer AD-associated protein signature, and integrated RNA–protein analysis revealed a smaller subset of coordinated molecular changes across transcriptomic and proteomic levels. These findings support the use of ONS cells as an accessible patient-derived model for investigating disease-associated molecular changes in AD and MCI and highlight candidate pathways and proteins for further validation.

## 5. Conclusion, limitations, and future directions

Here we report a proteomic characterisation of patient-derived ONS cells from AD, MCI and HC individuals using complementary fraction-specific and reconstructed bulk proteomics approaches. Subcellular fractionation identified disease-associated protein abundance changes in cytoplasmic and nuclear-enriched fractions, while reconstructed bulk proteomics identified a clearer AD-associated protein signature. Integration with our previous transcriptomic analysis of the same ONS cohort revealed partial RNA-protein concordance, suggesting that only a subset of disease-associated molecular changes was coordinated at both the transcript and protein level.

These findings support the use of ONS cells as an accessible patient-derived *in vitro* model for investigating molecular changes associated with AD and MCI. However, these findings should be interpreted in the context of several limitations, including the relatively small cohort size, individual variability between patient-derived cell lines, the exploratory nature of several candidate protein changes, and the immature neuroglial-like phenotype of ONS cells. Therefore, while they provide an accessible patient-derived model for investigating disease-associated molecular changes relevant to AD and MCI, the findings should not be interpreted as direct representations of mature brain cell pathology or advanced-stage neurodegeneration.

Although selected proteins showed supportive western blot findings or reproducible trends, further validation in larger independent cohorts will be required to confirm the robustness of these protein signatures. Given the number of altered proteins identified, only a subset could be assessed using targeted assays, selected according to statistical support, effect size and biological relevance. In addition, failure to reproduce individual proteomics findings by antibody-based validation methods does not necessarily indicate that the proteomics result is incorrect, as western blotting may be affected by differences in sensitivity, antibody specificity, epitope recognition, cross-reactivity, protein isoforms, post-translational modifications or fraction-specific protein detection [92–94]. Therefore, individual candidate proteins should be considered exploratory until confirmed in larger cohorts or using complementary validation approaches.

Future studies should also include larger independent cohorts, longitudinal sampling where possible, targeted proteomics, immunofluorescence validation of selected candidates and functional assays to determine whether the observed protein abundance and localisation changes contribute to AD-related cellular phenotypes. Together, these findings support the use of patient-derived ONS cells as an accessible *in vitro* model for investigating molecular changes associated with AD and MCI, while highlighting the potential of ONS cell-based proteomics to identify disease-associated protein signatures.

## 6. Materials and methods

### 6.1. Ethical requirements

Olfactory biopsies were performed with the approval of the Human Research Ethics Committees of Northern Savo Hospital District (permit number 536/2017) and QIMR Berghofer Medical Research Institute (P2197). Written informed consent was collected from all participants, with proxy consent obtained from family members of individuals with mild AD dementia. All research was conducted in accordance with the human research ethics guidelines of the National Health and Medical Research Council of Australia (NHMRC), QIMR Berghofer Medical Research Institute and the Northern Savo Hospital District.

### 6.2. Patients and nasal biopsy

Volunteers clinically diagnosed with MCI and mild AD dementia (CDR 1), together with cognitively healthy control subjects, were recruited via the Brain Research Unit, Department of Neurology, University of Eastern Finland, as previously described (Rantanen et al., 2022). Prior to study recruitment, diagnostic examinations had been carried out at the Brain Research Unit or at the Department of Neurology, Kuopio University Hospital (Lampinen, Fazaludeen, et al., 2022; Lampinen, Górová, et al., 2022). Olfactory mucosal biopsy collection and sample processing were performed according to established protocols (Féron et al., 1998; Murrell et al., 2005; Lampinen, Fazaludeen, et al., 2022; Rantanen et al., 2022). Briefly, olfactory mucosal tissue was obtained from the nasal septum and transported on ice in Dulbecco’s Modified Eagle Medium/Ham F-12 (DMEM/F12) (#11320033)-based growth medium supplemented with 10% heat-inactivated fetal bovine serum (FBS) (#10500064) and 1% penicillin-streptomycin solution (P/S) (#15140122) until processed in the laboratory. All reagents were obtained from Thermo Fisher Scientific. Primary olfactory mucosal cell cultures were established at the University of Eastern Finland using previously published methods (Féron et al., 1998; Lampinen, Fazaludeen, et al., 2022). For the proteomics analyses, 15 ONS cell lines were included: HC (*n* = 4), MCI (*n* = 6) and AD (*n* = 5). The demographic and clinical characteristics reported below refer to this proteomics cohort. The average ages of the AD, MCI and HC groups were approximately 70, 72 and 69 years, respectively. The AD group included three females and two males, the HC group included two females and two males and the MCI group included two females and four males. APOE genotyping showed that at least one APOE ε4 allele was present in 40% of patients with AD, 33% of participants with MCI and 50% of HC subjects (**Table S4**). APOE genotyping was performed as previously described (Huovinen et al., 2016). Olfactory function was assessed using the 12-odour Sniffin’ Sticks test, and participants were classified as normosmic, hyposmic or anosmic. Additional available ONS cell lines were used for western blot and Seahorse analyses. Western blot experiments included HC (*n* = 7), MCI (*n* = 8) and AD (*n* = 9) cell lines, while the Seahorse XF Cell Mito Stress Test included HC (*n* = 5), MCI (*n* = 6) and AD (*n* = 6), as described in Sections 6.15 and 6.16, respectively.

### 6.3. Generation of olfactory mucosa (OM)-derived neurospheres

OM cell lines derived from nasal biopsies were cultured as previously described [23]. Briefly, OM cells were expanded in DMEM/F12 culture medium supplemented with 10% FBS and 1% P/S until approximately 70% confluent. For neurosphere generation, cells were seeded at 12,000 cells/cm² into poly-L-lysine-coated T25 Nunc™ flasks (1 µg/cm², Sigma-Aldrich, #P4707) and cultured in sphere-inducing medium. This medium consisted of DMEM/F12 supplemented with 1% insulin-transferrin-selenium (ITS, Thermo Fisher Scientific, #41400-045), epidermal growth factor (EGF, 50 ng/mL, EDM Millipore, #GF144) and fibroblast growth factor-2 (FGF2, 25 ng/mL, EDM Millipore, #GF003-AF). Growth factors were replenished every two days. Typical passage number of OM cells before neurosphere induction was between 2 and 5.

### 6.4. Establishment of olfactory neurosphere-derived (ONS) cell cultures

Neurospheres detached or loosened from the culture surface after 2-7 days, depending on the cell line. Free-floating neurospheres were collected every second day together with the sphere-inducing culture medium, centrifuged at 100 × *g* for 5 min, and dissociated into single cells using TrypLE at 37°C for 10 min. Dissociated ONS cells were then re-seeded at 4,000–6,500 cells/cm^2^ into T75 Nunc^TM^ culture flasks (Nunc) and maintained in DMEM/F12 supplemented with 10% FBS until approximately 75% confluent, typically in 1–2 weeks. All the experiments were performed using ONS cells below passage 10.

### 6.5. ONS cell sample collection for omics analyses

Of the original six ONS cell lines per diagnostic group, 15 lines met the sample quantity and quality requirements for subcellular fractionation and proteomics analysis and were included in the omics analyses: HC (*n* = 4), MCI (*n* = 6) and AD (*n* = 5). Cells were seeded at 4,000–6,500 cells/cm^2^ into Corning^TM^ TC-treated culture dishes (Corning, #430599) and cultured in DMEM/F12 medium supplemented with 10% FBS. Cells were expanded until approximately 75% confluent, then manually detached using a cell scraper and 1 mL of ice-cold PBS to minimise proteolysis. Cells were centrifuged at 800 × *g* for 5 min at +4°C, and pellets containing approximately one million cells per line were frozen and stored at -80°C until subcellular fractionation. Proteomics analysis was subsequently performed by LC-MS/MS.

### 6.6. RNA extraction, cDNA synthesis and quantitative real-time PCR (qRT-PCR)

Total RNA was extracted from ONS cells using TRIzol™ reagent (Life Technologies) and the Direct-zol RNA Miniprep Kit (Zymo Research, #R2052) according to the manufacturer’s instructions. Briefly, cells were lysed in TRIzol™, mixed with ethanol and transferred to Zymo-Spin™ columns for RNA binding. Columns were washed and treated with DNase I on-column to remove genomic DNA contamination before RNA was eluted in DNase/RNase-free water. RNA concentration and purity were assessed using a NanoDrop™ spectrophotometer.

For qRT-PCR analysis, 150 ng of total RNA was converted to cDNA using the SensiFAST™ cDNA Synthesis Kit (Bioline, #BIO-65054) according to the manufacturer’s instructions. cDNA was diluted 1:10 and qRT-PCR was performed using SensiFAST™ SYBR® Lo-ROX master mix with gene-specific primers on a QuantStudio™ 5 Real-Time PCR System (Thermo Fisher Scientific). Reactions were run in technical triplicates using the following cycling conditions: 95°C for 2 min, followed by 40 cycles of 95°C for 5 s and 60°C for 20 s.

Ct values were normalised to the endogenous control 18S, and relative gene expression was calculated using the 2^−ΔCt method. Technical triplicates were averaged for each biological sample before statistical analysis. Primer sequences are listed in Table S5.

### 6.7. Immunofluorescence of olfactory neurosphere-derived cells

ONS cells were cultured on 13 mm plastic coverslips (Sarstedt) in DMEM/F12 medium containing 10% FBS and 1% penicillin/streptomycin. Cells were fixed with either 4% paraformaldehyde for 15 min or ice-cold methanol for 5 min, followed by PBS washes. PFA-fixed samples were permeabilised with PBS containing 0.5% Triton X-100. Samples were blocked for 2 h at room temperature in PBS containing 2% bovine serum albumin, 2% goat serum and 0.1% Triton X-100. Primary antibodies were diluted in blocking solution and incubated overnight at 4°C. The following day, cells were washed three times with PBS containing 0.1% Triton X-100 and incubated with secondary antibodies for 2 h at room temperature in the dark. Nuclei were counterstained with Hoechst. Primary and secondary antibodies are listed in Table S6. Fluorescence images were captured using a confocal laser scanning microscope LSM-780 (Carl Zeiss) and processed using Zeiss ZEN software.

### 6.8. Subcellular fractionation

Approximately one million propagated ONS cells were pelleted and stored at -80°C until subcellular fractionation. Cell pellets were thawed on ice and resuspended in chilled hypotonic lysis buffer containing 10 mM Tris pH 7.5, 2 mM EDTA, 25 mM sodium fluoride and 1 × protease inhibitor cocktail. Samples were incubated on ice for 15 min, followed by mechanical disruption using a 24G syringe. Nuclei were pelleted by centrifugation at 1,000 × *g* for 10 min at 4°C, and the supernatant containing the cytoplasmic fraction was collected. The cytoplasmic fraction was centrifuged again to remove residual nuclei or intact cells. The nuclear pellet was washed with PBS and homogenised in a small volume of PBS. Protein concentration in each nuclear and cytoplasmic fraction was determined using the BCA assay. Although protein yield varied between samples and fractions, sufficient material was obtained for proteomic preparation. For samples containing less than 20 µg protein, the entire sample was digested and trypsin was used at a 1:20 ratio. The efficiency of subcellular fractionation was initially assessed in test samples by western blotting for the nuclear marker DDX5 and cytoplasmic marker HSP90, with actin used as a loading control (data not shown). Although western blotting was not performed for every study sample, the proteomics data detected known nuclear and cytoplasmic marker proteins in the expected fractions, supporting successful enrichment of the respective compartments (**Fig. S10**). Mitochondrial and endoplasmic reticulum (ER)-associated proteins were detected predominantly in the cytoplasmic fraction, as expected (**Fig. S10**).

### 6.9. Protein precipitation and digestion

Immediately following fractionation, three volumes of methanol were added to each fraction and samples were incubated overnight at -20°C. Precipitated proteins were pelleted by centrifugation at 16,000 × *g* for 30 min at 4°C, gently washed with 50 mM triethylammonium bicarbonate (TEAB), and resuspended in TEAB. Protein concentration was determined using a BCA assay. For each sample, 20 µg of protein was prepared in TEAB at a final concentration of 1 µg/µL. Sodium deoxycholate was added to a final concentration of 1% to support solubilisation of membrane-associated proteins during digestion. A modified in-solution digestion protocol was performed using the BRAVO AssayMap liquid handling platform. Briefly, samples were reduced with 5mM dithiothreitol (DTT) for 30 min at 37°C and alkylated with 20 mM 2-Iodoacetamide (IAA) for 25 min. Samples were then diluted 1:10 with TEAB, and 0.7 µg trypsin was added for overnight digestion at 37°C. Digestion was stopped by adding trifluoroacetic acid (TFA) to a final concentration of 0.5%, which also precipitated sodium deoxycholate. Samples were centrifuged at 5,000 × *g* for 30 min, and the resulting supernatants were desalted using AssayMap C18 cartridges according to the standard protocol. Eluted peptides were resuspended in 0.5% TFA, and stored at -20°C until LC-MS/MS analysis.

### 6.10. Proteomics acquisition and protein identification

Peptides were separated using a Thermo U3000 nanoHPLC system and analysed on a Thermo Q Exactive Plus Orbitrap mass spectrometer. Peptides were loaded onto a C-18 trap column and resolved using a 50 cm EasySpray C-18 analytical column (Thermo Fisher, #160454 and ES803A). Mobile phase A consisted of 0.1% formic acid in water, and mobile phase B consisted of 80% acetonitrile with 0.1% formic acid. Samples were loaded in 3% mobile phase B at 10 μL/min, with an injection volume of 4 µL equivalent to approximately 1 µg peptide. Peptides were separated at a flow rate of 250 nL/min using a multi-step gradient from 3% to 6% B over 1 min, 6% to 30% B over 60 min, 30% to 50% B over 12 min, 50% to 95% B over 1 min. The column was then maintained at 95% B for 12 min before re-equilibration to 3% B. The mass spectrometer was operated in positive ion data-dependent acquisition mode using an EasySpray source. Full MS scans were acquired over a mass range of 350–1400 m/z at 70,000 resolution, with an AGC target of 3 × 10⁶ and a maximum injection time of 100 ms. MS/MS fragmentation was performed on the top 10 precursor ions, excluding 1+ charged and >7+ charged precursors. The dynamic exclusion window was 30 s, precursor isolation width was 1.4 m/z, and normalised collision energy was 27. MS/MS scans were acquired at 17,500 resolution, with an AGC target of 5 × 10⁵ and a maximum injection time of 50 ms. Mass lock was set to “Best”. Raw files were processed using MaxQuant for protein identification and quantification. Spectra were searched against the reviewed human Swiss-Prot proteome database, version 2021_03. False discovery rates were set at 5% at both peptide and protein levels. Methionine oxidation and N-terminal acetylation were included as variable modifications. Protein identification required a minimum of two unique peptides, and quantification was performed using the razor peptide method. Match between runs was enabled, and all other parameters were kept at default settings.

### 6.11. Proteomics data analysis

Proteomics data were analysed using the LFQ-Analyst web platform. Protein abundance was quantified using label-free quantification (LFQ) intensity values, which provide relative abundance estimates suitable for comparison between samples and diagnostic groups, but do not represent absolute protein concentrations. Protein log_2_ fold changes, *p*-values and imputed intensity values were obtained from LFQ-Analyst after preprocessing using default statistical parameters. Differential protein abundance analysis was performed using protein-wise linear models with empirical Bayes moderation implemented through the Bioconductor package limma, as described previously (Shah et al., 2020). Pairwise comparisons were performed between HC, MCI and AD samples within the cytoplasmic and nuclear fractions. Differentially abundant proteins (DEPs) were identified using FDR-adjusted *p*-values. Proteins passing a nominal *p* < 0.01 threshold were considered exploratory candidate DEPs and were used for visualisation and interpretation where indicated. Gene ontology enrichment analysis was performed using the PANTHER overrepresentation test at geneontology.org. Proteins altered at *p* < 0.05 in each comparison were used as input, and enriched terms were considered significant at FDR < 0.05. PANTHER GO-Slim Biological Process and PANTHER Protein Class annotations were analysed using Fisher’s exact test.

### 6.12. Protein quality control analysis

Data quality assessment was conducted before biological interpretation to evaluate protein coverage, sample consistency and overall data consistency [95]. Protein intensity distributions were assessed before and after normalisation and showed broadly comparable distributions across samples, supporting the suitability of the normalised dataset for downstream analysis (**Fig. S1A, B**). Protein coverage was then evaluated separately in the cytoplasmic and nuclear fractions. Cytoplasmic samples contained approximately 2,400–2,700 proteins per sample, while nuclear samples contained approximately 2,100–2,500 proteins per sample, consistent with the lower complexity of the nuclear fraction (**Fig. S1C**). Sample-level variability was further assessed using coefficient of variation plots (**Fig. 1D**) and principal component analysis. Coefficient of variation distributions showed moderate variability within diagnostic groups, as expected for patient-derived human cell lines, with median CV values ranging from approximately 26-35% across cytoplasmic and nuclear sample groups (**Fig. 1D**). PCA of the cytoplasmic and nuclear fractions showed partial separation of AD ONS lines from HC and MCI samples, while HC and MCI samples showed greater overlap and clustered more closely together in both fractions (**Fig. 2A, B**).

### 6.13. Reconstructed bulk proteomics

For reconstructed bulk proteomics analysis, cytoplasmic and nuclear-enriched fraction data from the same ONS cell lines used in the subcellular proteomics workflow were integrated at the MaxQuant processing stage. The final reconstructed bulk dataset included HC (*n* = 4), MCI (*n* = 6) and AD (*n* = 5) ONS cell lines. To approximate a bulk proteomics dataset, mass spectrometry data from the cytoplasmic and nuclear-enriched fractions were re-aligned to improve protein identification and quantification across the combined dataset. Peptide intensity distributions were assessed before and after log transformation and normalisation to confirm comparable intensity distributions across samples (**Fig. S5**). Protein abundance was quantified using label-free quantification (LFQ) intensity values, which provide relative abundance estimates suitable for comparison between samples and diagnostic groups, but do not represent absolute protein concentrations. LFQ intensity data were filtered to exclude proteins missing in at least three samples, resulting in a final dataset of 2,490 proteins.

Differential protein abundance between diagnostic groups was assessed for AD vs HC, AD vs MCI and MCI vs HC comparisons using limma in R with moderated *t*-tests. Adjusted *p*-values were calculated using the Benjamini–Hochberg method to control the false discovery rate. Proteins passing FDR correction were considered significantly differentially abundant.

Gene Ontology (GO) enrichment analysis was performed using the clusterProfiler package in R on proteins identified as significantly differentially abundant after Benjamini–Hochberg correction. As only the AD vs HC comparison yielded a sufficient number of significant proteins, GO enrichment analysis was performed for this comparison.

### 6.14. Transcriptomic-proteomics correlation analysis

Transcriptomic and proteomic datasets were integrated by matching genes detected in the reconstructed bulk proteomics dataset and the previously reported RNA-seq dataset from the same ONS cohort [23]. Only genes present in both datasets were retained. For genes with multiple isoforms or transcripts, the transcript with the highest median expression was selected. Prior to correlation analysis, both datasets were log-transformed and mean-centred using the limma package in R. For each matched RNA–protein pair, normality was assessed using the Shapiro– Wilk test. Matched pairs that met the normality assumption (*p* > 0.05) were analysed using Pearson’s correlation to evaluate the linear relationship between transcript abundance and protein intensity. Pearson’s correlation coefficients (*r*) and corresponding 95% confidence intervals were calculated for each analysed pair.

### 6.15. Western blot

ONS cells from HC (*n* = 7), MCI (*n* = 8) and AD (*n* = 9) individuals were used for western blot analyses. Cells were rinsed with ice-cold PBS and lysed in radioimmunoprecipitation assay buffer (RIPA) (Merck, #20-188) supplemented with protease and phosphatase inhibitor cocktails (Roche #11873580001 and #04906845001). Lysates were incubated on ice for 10 min and then centrifuged at 2060 × g for 15 min at 4°C. Supernatants were collected, and protein concentrations were determined using the Pierce™ BCA Protein Assay Kit (Thermo Fisher Scientific, #23225) according to the manufacturer’s instructions. Equal amounts of protein lysate (20 μg) were separated by SDS-PAGE using an Invitrogen Mini Gel Tank system (Thermo Fisher Scientific, #A25977) with Bolt^TM^ 4–12% Bis-Tris Plus polyacrylamide gels and 1 × MES SDS running buffer (Thermo Fisher Scientific). Proteins were transferred onto polyvinylidene difluoride (PVDF) membranes (Invitrogen #IB34002) using the iBlot™ 3 Dry Blotting System (Thermo Fisher Scientific). Membranes were blocked for 1 h at RT in 5% bovine serum albumin (BSA, Sigma Aldrich, #A7906-100G) prepared in Tris-buffered saline containing 0.1% Tween-20 (TBS-T). Membranes were then incubated overnight at 4°C with the following primary antibodies diluted in 5% BSA/TBS-T: anti-S100A4 (1:500, Invitrogen, #MA5-32347), anti-LC3B (1:1000, Invitrogen, #PA1-46286), anti-NDUFS4 (1:500, Abcam, #ab137064), anti-PSEN1 (1:1000, Cell Signaling Technology, #5643S) and anti-GAPDH (1:10,000, Cell Signaling Technology, #2118S). The following day, membranes were washed with TBS-T and incubated for 1 h at RT with horseradish peroxidase-conjugated secondary antibody (1:10,000, Cell Signaling Technology, #7074S) diluted in 5% BSA in TBS-T. After washing with TBS-T, the immunoreactive bands were visualised using SuperSignal™ West Pico Plus Chemiluminescent Substrate (Thermo Fisher Scientific #34580) and captured using an iBright™ CL1500 Imaging System (Thermo Fisher Scientific). Band intensities were quantified using FIJI ImageJ Version 1.54p. Target protein abundance was normalised to GAPDH, which served as the loading control, and expressed relative to control samples.

### 6.16. Seahorse XF Cell Mito Stress Test

The Seahorse XF Cell Mito Stress Test was performed to assess mitochondrial respiration in HC (*n* = 5), MCI (*n* = 6) and AD (*n* = 6) ONS cells according to the manufacturer’s instructions. ONS cells were seeded onto Seahorse XFe96 cell culture microplates at a density of 15,000 cells per well and cultured for 48 h in DMEM/F12 medium supplemented with 10% FBS and 1% P/S. Five technical replicates were included for each cell line. After 48 h, cells were washed once with Seahorse XF DMEM medium, pH 7.4 (Agilent #103575-100), supplemented with 25 mM D-glucose (Sigma-Aldrich #G8769), 2 mM sodium pyruvate (Gibco #11360-070) and 2 mM GlutaMAX (Gibco #35050-061). Cells were then incubated at 37°C in a CO₂-free environment for 1 h before analysis. Oxygen consumption rate (OCR) was measured using the Seahorse XFe96 Analyzer (Agilent). Baseline OCR was recorded before sequential injection of oligomycin (ATP synthase inhibitor, Sigma-Aldrich #75351), 1 µM FCCP (oxidative phosphorylation uncoupler, Sigma-Aldrich, #C2920), and a mix of rotenone and antimycin A (Complex I and III inhibitors, 1 µM each, Sigma-Aldrich, #R8875 and #A8674). OCR values were normalised to total protein content. Respiratory parameters were calculated using Seahorse Wave software (Agilent). Basal respiration was calculated as the last OCR measurement before oligomycin injection minus non-mitochondrial respiration, ATP production was calculated as basal OCR minus OCR following oligomycin treatment, maximal respiration as the maximum OCR following FCCP treatment minus non-mitochondrial respiration, spare respiratory capacity as maximal respiration minus basal respiration, proton leak as OCR following oligomycin treatment minus non-mitochondrial respiration, and coupling efficiency as [(basal OCR − OCR after oligomycin)/basal OCR] × 100. Non-mitochondrial oxygen consumption was determined from OCR following rotenone/antimycin A treatment. Wells showing negative OCR values or visible cell detachment around the sensor area were excluded as technical failures. Outliers among biological replicate values were assessed using Grubbs’ test (α = 0.05). One HC donor value was identified as an outlier for proton leak and excluded from the primary analysis, no other biological data points were excluded. For each biological cell line, values from the remaining technical replicate wells were averaged to generate a single value for statistical analysis. Statistical comparisons were performed using biological cell lines as the independent observations.

### 6.17. Statistical analysis

GraphPad Prism software version 10.3.0 (GraphPad Software Inc., San Diego, CA, USA) was used to produce graphs and statistical analyses. Data distribution was assessed for normality before statistical testing using GraphPad Prism normality tests, including the Shapiro-Wilk test. Comparisons between two groups were performed using two-tailed unpaired *t*-tests with Welch’s correction. Comparisons between three groups were performed using one-way analysis of variance (ANOVA) followed by Tukey’s post hoc tests. Data are presented as mean ± SEM unless otherwise stated, and *p* ˂ 0.05 was considered statistically significant. Statistical significance is indicated in the figures as \**p* < 0.05, \*\**p* < 0.01, \*\*\**p* < 0.001, \*\*\*\**p* < 0.0001. Potential outliers were assessed before statistical testing using the GraphPad Outlier calculator based on Grubbs’ test with α = 0.05. Outliers identified by this method were excluded from the downstream analysis. The number of biological replicates used for each experiment is indicated in the relevant figure legends.

### 6.19. Data availability

The mass spectrometry proteomics data have been deposited in the ProteomeXchange Consortium via the PRIDE partner repository with the dataset identifier PXD082268.

## Reviewer access details

Log in to the PRIDE website using the following details:

## Project accession

PXD082268

## Token

HSBRVkT92L7k

## Acknowledgements

The authors acknowledge the technical assistance provided by the Proteomics Facility at the Translational Research Institute (TRI), Brisbane. The authors thank Amali Fernando for assistance with western blot experiments. The authors also thank research nurses Pirkko Kanerva and Hanna Kämppi for their invaluable support with participant contact and study coordination, demographic and clinical interviews, cognitive and other clinical assessments, blood sample collection, and preparation of the clinical data records used for subsequent analyses.

## Ethics Approval

The study was conducted in accordance with the Declaration of Helsinki and approved by the Human Research Ethics Committees of Northern Savo Hospital District (permit no. 536/2017) and QIMR Berghofer Medical Research Institute (P2197). All research was conducted in accordance with applicable ethical guidelines for human research.

## Consent to Participate

Informed consent was obtained from all participants involved in the study.

