## Supplementary material for "Alzheimer’s Disease-Associated Proteomic Changes in Olfactory Stem Cells Show Pathway-Level Transcriptomic Overlap": Supplementary material_Rantanen et al.pdf

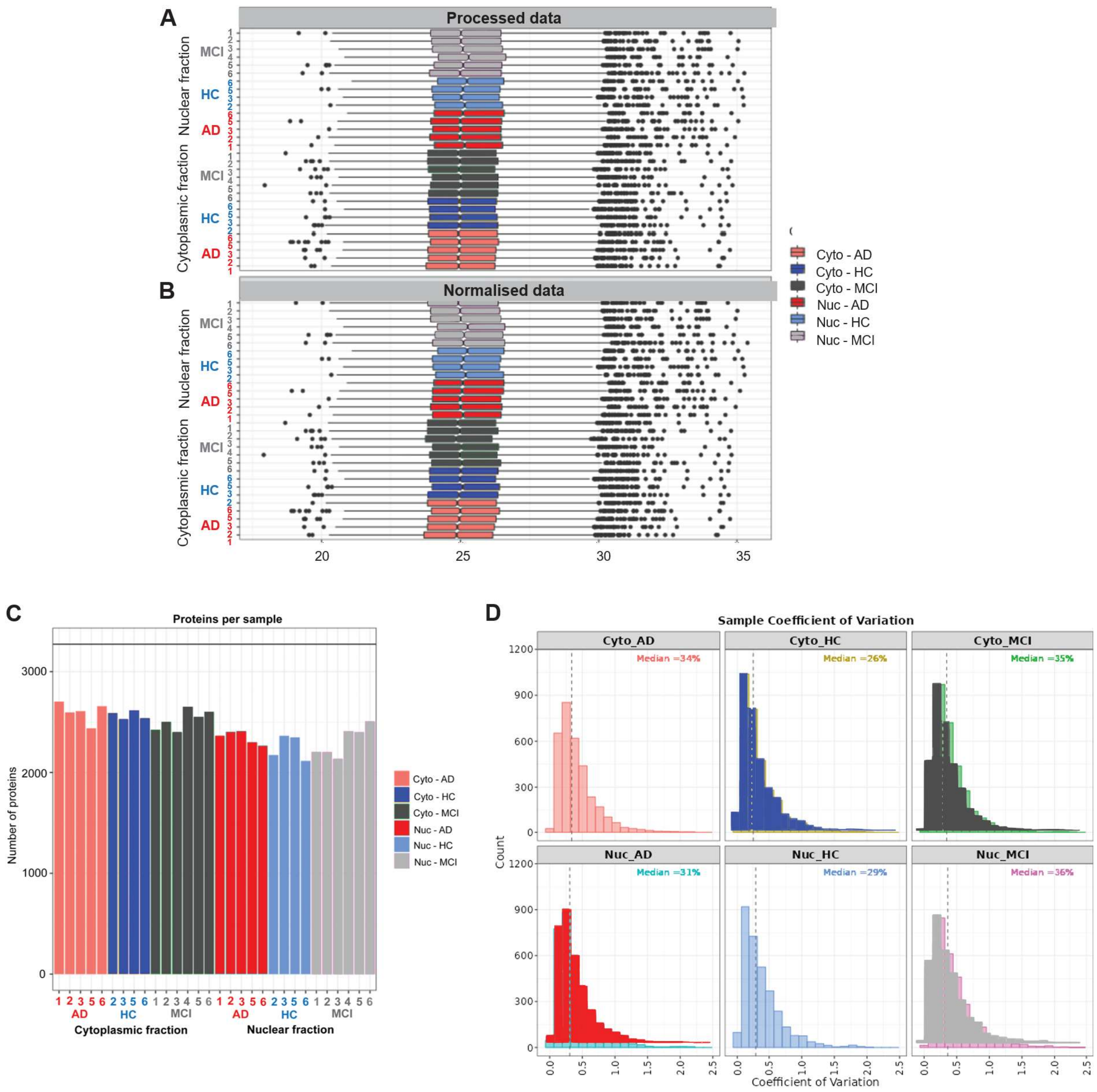


**Fig. S1. Proteomics data quality control.** (**A**) Pre- and (**B**) post-normalised protein intensity distributions across cytoplasmic and nuclear-enriched fractions. (**C**) Number of proteins identified per sample in cytoplasmic and nuclear-enriched fractions. (**D)** Sample coefficient of variation distributions for AD, HC and MCI ONS lines in cytoplasmic and nuclear-enriched fractions. All plots were generated using LFQ-Analyst.

**Table S1. Selected metabolism-related and other candidate differentially abundant proteins identified across AD, MCI and HC ONS cell comparisons in nuclear-enriched (Nuc) and cytoplasmic (Cyto) fractions.** Most proteins were selected using a nominal threshold of $p < 0.01$. Additional metabolism-related proteins with $p < 0.05$, including selected NADH dehydrogenase subunits, were included due to their biological relevance to the Complex I-related proteomic pattern. $Log_2FC$, $p$-values and FDR values were determined using LFQ-Analyst. Fraction labels indicate whether the change was detected in the nuclear-enriched or cytoplasmic fraction.

| | AD vs HC | | | AD vs MCI | | | MCI vs HC | | |
|---|---|---|---|---|---|---|---|---|---|
| **Name** | **log₂FC** | ***p*-values** | **FDR** | **log₂FC** | ***p*-values** | **FDR** | **log₂FC** | ***p*-values** | **FDR** |
| **Mitochondrial- and metabolism-associated proteins** | | | | | | | | | |
| Complex I/NADH dehydrogenase-associated subunits | | | | | | | | | |
| NDUFS4 | -1.90 (Nuc) | 0.004 (Nuc) | 0.782 (Nuc) | -3.58 (Nuc) | 8.67 x $10^{-6}$ (Nuc) | **0.009 (Nuc)** | 1.68 (Nuc) | 0.007 (Nuc) | 0.485 (Nuc) |
| NDUFS6 | 1.36 (Nuc) | 0.005 (Nuc) | 0.803 (Nuc) | 0.71 (Cyto) | 0.050 (Cyto) | 0.912 (Cyto) | | | |
| NDUFS7 | | | | -2.02 (Nuc) | 0.003 (Nuc) | 0.527 (Nuc) | -1.55 (Nuc) | 0.015 (Nuc) | 0.527 (Nuc) |
| NDUFS8 | -0.84 (Cyto) | 0.029 (Cyto) | 0.434 (Cyto) | | | | | | |
| NDUFS1 | -0.46 (Nuc) | 0.022 (Nuc) | 0.902 (Nuc) | | | | | | |
| NDUFA13 | -0.64 (Nuc) | 0.023 (Nuc) | 0.906 (Nuc) | | | | | | |
| NDUFA5 | 0.48 (Nuc) | 0.058 (Nuc) | 0.939 (Nuc) | 1.33 (Cyto) | 0.030 (Cyto) | 0.850 (Cyto) | 0.52 (Nuc); -1.40 (Cyto) | 0.036 (Nuc); 0.031 (Cyto) | 0.737 (Nuc); 0.859 (Cyto) |
| NDUFA6 | | | | -1.41 (Cyto) | 0.030 (Cyto) | 0.859 (Cyto) | | | |
| NDUFAB1 | | | | | | | -1.54 (Cyto) | 0.045 (Cyto) | 0.943 (Cyto) |
| NDUFAF4 | -0.99 (Cyto) | 0.038 (Cyto) | 0.475 (Cyto) | | | | | | |
| Glycolysis-associated proteins | | | | | | | | | |
| ENO1 | 1.86 (Nuc) | 6.85 × $10^{-5}$ (Nuc) | 0.422 (Nuc) | 1.81 (Nuc) | 5.76 × $10^{-5}$ (Nuc) | 0.063 (Nuc) | | | |
| TPI1 | 1.72 (Nuc) | 0.0005 (Nuc) | 0.444 (Nuc) | 1.57 (Nuc) | 0.0007 (Nuc) | 0.318 (Nuc) | | | |
| GAPDH | 0.55 (Nuc) | 0.010 (Nuc) | 0.857 (Nuc) | 0.73 (Nuc) | 0.0009 (Nuc) | 0.358 (Nuc) | | | |
| ALDOA | 0.76 (Nuc) | 0.002 (Nuc) | 0.678 (Nuc) | 0.66 (Nuc) | 0.004 (Nuc) | 0.571 (Nuc) | | | |
| HK2 | | | | 1.36 (Nuc) | 0.017 (Nuc) | 0.762 (Nuc) | -1.42 (Nuc) | 0.013 (Nuc) | 0.556 (Nuc) |
| PGAM1 | 1.62 (Nuc) | 0.0007 (Nuc) | 0.493 (Nuc) | 1.44 (Nuc) | 0.001 (Nuc) | 0.397 (Nuc) | | | |
| Complex IV-associated subunits | | | | | | | | | |
| COX5A | 0.53 (Nuc) | 0.028 (Nuc) | 0.913 (Nuc) | 1.76 (Cyto) | 0.041 (Cyto) | 0.893 (Cyto) | 0.47 (Nuc) | 0.040 (Nuc) | 0.757 (Nuc) |

| | | | | | | | | | |
|---|---|---|---|---|---|---|---|---|---|
| COX5B | 1.94 (Nuc); 2.18 (Cyto) | 0.0004 (Nuc); 0.019 (Cyto) | 0.439 (Nuc); 0.374 (Cyto) | 1.79 (Cyto) | 0.031 (Cyto) | 0.864 (Cyto) | 1.22 (Nuc) | 0.009 (Nuc) | 0.514 (Nuc) |
| COX6B1 | 2.13 (Cyto) | 0.001 (Cyto) | **0.047 (Cyto)** | 1.26 (Cyto) | 0.021 (Cyto) | 0.802 (Cyto) | | | |
| COX7C | 3.65 (Cyto) | 0.0002 (Cyto) | **0.003 (Cyto)** | | | | 2.58 (Cyto) | 0.002 (Cyto) | 0.098 (Cyto) |
| ATP synthase/proton transport-associated proteins | | | | | | | | | |
| ATP5J | 1.11 (Nuc); 3.39 (Cyto) | 0.012 (Nuc); 0.0007 (Cyto) | **0.017 (Nuc)** | 2.13 (Cyto) | 0.009 (Cyto) | 0.611 (Cyto) | | | |
| ATP6V1F | 1.92 (Cyto) | $7.55 \times 10^{-5}$ (Cyto) | **0.001 (Cyto)** | 1.07 (Cyto) | 0.004 (Cyto) | 0.421 (Cyto) | 0.853 (Cyto) | 0.024 (Cyto) | 0.461 (Cyto) |
| ATP6V0A1 | -1.73 (Nuc) | 0.007 (Nuc) | 0.827 (Nuc) | | | | -1.34 (Nuc) | 0.023 (Nuc) | 0.643 (Nuc) |
| ATP6V0D1 | -0.68 (Nuc) | 0.0008 (Nuc) | 0.520 (Nuc) | | | | -0.38 (Nuc) | 0.028 (Nuc) | 0.684 (Nuc) |
| ATP6V1E1 | | | | 1.12 (Nuc) | 0.0416 (Nuc) | 0.860 (Nuc) | -1.35 (Nuc) | 0.0166 (Nuc) | 0.581 (Nuc) |
| ATP6V1A | 0.65 (Nuc) | 0.030 (Nuc) | 0.917 (Nuc) | | | | | | |
| ATP6V1B2 | | | | | | | 0.32 (Nuc) | 0.046 (Nuc) | 0.779 (Nuc) |
| Ion transport/membrane potential-related proteins | | | | | | | | | |
| ATP1B3 | | | | | | | 0.55 (Cyto) | 0.023 (Cyto) | 0.451 (Cyto) |
| MCU | -1.20 (Nuc) | 0.0003 (Nuc) | 0.433 (Nuc) | | | | -0.742 (Nuc) | 0.008 (Nuc) | 0.503 (Nuc) |
| Mitochondrial ribosomal/translation-associated proteins | | | | | | | | | |
| MRPL1 | 0.94 (Cyto) | 0.019 (Cyto) | 0.366 (Cyto) | | | | 0.76 (Cyto) | 0.042 (Cyto) | 0.588 (Cyto) |
| MRPL3 | 1.11 (Cyto) | 0.034 (Cyto) | 0.453 (Cyto) | | | | | | |
| MRPL12 | 1.36 (Cyto) | 0.028 (Cyto) | 0.425 (Cyto) | 1.14 (Cyto) | 0.040 (Cyto) | 0.892 (Cyto) | | | |
| MRPL21 | -1.11 (Nuc) | 0.0003 (Nuc) | 0.433 (Nuc) | | | | -1.12 (Nuc) | 0.0002 (Nuc) | **0.0371 (Nuc)** |
| MRPL28 | 1.69 (Cyto) | 0.0003 (Cyto) | **0.005 (Cyto)** | | | | 1.25 (Cyto) | 0.003 (Cyto) | 0.106 (Cyto) |
| MRPL34 | 1.10 (Cyto) | 0.002 (Cyto) | 0.064 (Cyto) | | | | 0.69 (Cyto) | 0.025 (Cyto) | 0.465 (Cyto) |
| MRLP45 | | | | 4.07 (Cyto) | 0.012 (Cyto) | 0.681 (Cyto) | -4.57 (Cyto) | 0.009 (Cyto) | 0.272 (Cyto) |

| | | | | | | | | | |
|---|---|---|---|---|---|---|---|---|---|
| MRPL47 | | | | | | | 1.17 (Cyto) | 0.008 (Cyto) | 0.256 (Cyto) |
| Mitochondrial protein import / transport-associated proteins | | | | | | | | | |
| TIMM9 | 0.85 (Nuc) | 0.004 (Nuc) | 0.769 (Nuc) | | | | 0.77 (Nuc) | 0.005 (Nuc) | 0.429 (Nuc) |
| TIMM44 | -0.57 (Nuc) | 0.014 (Nuc) | 0.879 (Nuc) | | | | | | |
| TIMM8A | 1.91 (Nuc) | 0.005 (Nuc) | 0.801 (Nuc) | | | | 2.12 (Nuc) | 0.002 (Nuc) | 0.268 (Nuc) |
| MTX2 | -0.98 (Nuc) | 0.0002 (Nuc) | 0.429 (Nuc) | | | | -0.62 (Nuc) | 0.005 (Nuc) | 0.43 (Nuc) |
| Mitochondrial intermediary metabolism / TCA / redox-associated proteins | | | | | | | | | |
| IDH2 | -0.55 (Nuc) | 0.002 (Nuc) | 0.690 (Nuc) | -0.34 (Nuc) | 0.030 (Nuc) | 0.830 (Nuc) | | | |
| OGDH | -0.498 (Nuc) | 0.012 (Nuc) | 0.870 (Nuc) | -0.41 (Nuc) | 0.026 (Nuc) | 0.816 (Nuc) | | | |
| DLD | 1.15 (Nuc) | 0.032 (Nuc) | 0.920 (Nuc) | 0.98 (Nuc) | 0.053 (Nuc) | 0.879 (Nuc) | | | |
| ALDH2 | 0.99 (Nuc) | 0.005 (Nuc) | 0.799 (Nuc) | | | | | | |
| Mitochondrial metabolite transporters | | | | | | | | | |
| SLC25A13 | -0.67 (Nuc) | 0.004 (Nuc) | 0.764 (Nuc) | -0.73 (Nuc) | 0.001 (Nuc) | 0.434 (Nuc) | | | |
| SLC25A20 | -1.25 (Cyto) | 0.006 (Cyto) | 0.196 (Cyto) | | | | -1.27 (Cyto) | 0.005 (Cyto) | 0.170 (Cyto) |
| Glucose / amino acid / lipid metabolism-associated proteins | | | | | | | | | |
| SLC2A1 | | | | -0.49 (Nuc) | 0.019 (Nuc) | 0.777 (Nuc) | | | |
| AGK | -0.90 (Nuc) | 0.0004 (Nuc) | 0.442 (Nuc) | | | | -0.55 (Nuc) | 0.011 (Nuc) | 0.538 (Nuc) |
| PHGDH | | | | 1.19 (Cyto) | 0.0019 (Cyto) | 0.301 (Cyto) | | | |
| Other metabolism- or stress-associated proteins | | | | | | | | | |
| DDAH2 | 1.62 (Cyto) | 0.0002 (Cyto) | **0.003 (Cyto)** | | | | 1.14 (Cyto) | 0.003 (Cyto) | 0.108 (Cyto) |
| BCL2L13 | -1.43 (Nuc); -1.75 (Cyto) | $6.11 \times 10^{-5}$ (Nuc); 0.006 (Cyto) | 0.421 (Nuc); 0.180 (Cyto) | -0.56 (Nuc); -1.31 (Cyto) | 0.038 (Nuc); 0.017 (Cyto) | 0.852 (Nuc); 0.768 (Cyto) | -0.86 (Nuc) | 0.003 (Nuc) | 0.373 (Nuc) |
| **Other most significantly altered DEPs** | | | | | | | | | |
| GGCX | -0.76 (Nuc) | $1.18 \times 10^{-5}$ (Nuc) | 0.387 (Nuc) | -0.79 (Nuc) | $5.68 \times 10^{-6}$ (Nuc) | **0.009 (Nuc)** | | | |
| NID1 | | | | -3.35 | $2.23 \times 10^{-5}$ | **0.033** | 2.24 | 0.001 (Nuc) | 0.172 |

| | | | | | | | | | |
|---|---|---|---|---|---|---|---|---|---|
| | | | | (Nuc) | (Nuc) | **(Nuc)** | (Nuc) | | (Nuc) |
| EIF5B | 0.98 (Nuc) | 0.0002 (Nuc) | 0.432 (Nuc) | 1.17 (Nuc) | $3.14 \times 10^{-5}$ (Nuc) | **0.043 (Nuc)** | | | |
| CNTNAP1 | 0.83 (Nuc); 1.10 (Cyto) | 0.0002 (Nuc) and 0.001 (Cyto) | 0.429 (Nuc); **0.039 (Cyto)** | | | | 0.87 (Nuc); 0.61 (Cyto) | 0.0001 (Nuc); 0.036 (Cyto) | **0.018 (Nuc);** 0.549 (Cyto) |
| SCRIB | -1.27 (Nuc); -0.97 (Cyto) | 0.0003 (Nuc); 0.0002 (Cyto) | 0.432 (Nuc); **0.003 (Cyto)** | -0.39 (Cyto) | 0.048 (Cyto) | 0.908 (Cyto) | -1.20 (Nuc); -0.58 (Cyto) | 0.0003 (Nuc); 0.009 (Cyto) | **0.048 (Nuc);** 0.269 (Cyto) |
| PNPLA6 | -1.41 (Nuc) | $5.14 \times 10^{-5}$ (Nuc) | 0.419 (Nuc) | | | | -1.13 (Nuc) | 0.0003 (Nuc) | **0.049 (Nuc)** |
| CASK | -0.51 (Nuc) | 0.011 (Nuc) | 0.865 (Nuc) | 0.47 (Nuc) | 0.014 (Nuc) | 0.732 (Nuc) | -0.98 (Nuc) | $3.49 \times 10^{-5}$ (Nuc) | **0.004 (Nuc)** |
| FRMD6 | 1.06 (Nuc); 0.62 (Cyto) | $1.40 \times 10^{-5}$ (Nuc); 0.046 (Cyto) | 0.387 (Nuc); 0.510 (Cyto) | 0.60 (Nuc) | 0.0018 (Nuc) | 0.464 (Nuc) | 0.46 (Nuc) | 0.0108 (Nuc) | 0.534 (Nuc) |
| PPIA | 1.02 (Nuc) | $2.65 \times 10^{-5}$ (Nuc) | 0.409 (Nuc) | 0.67 (Nuc) | 0.001 (Nuc) | 0.371 (Nuc) | 0.36 (Nuc) | 0.047 (Nuc) | 0.783 (Nuc) |
| YWHAZ | 0.95 (Nuc) | $2.82 \times 10^{-5}$ (Nuc) | 0.410 (Nuc) | 0.697 (Nuc) | 0.0004 (Nuc) | 0.241 (Nuc) | | | |
| YWHAB | 1.03 (Nuc) | $3.50 \times 10^{-5}$ (Nuc) | 0.414 (Nuc) | 0.72 (Nuc) | 0.0007 (Nuc) | 0.322 (Nuc) | | | |
| AKAP12 | 1.16 (Nuc) | 0.0009 (Nuc) | 0.536 (Nuc) | -0.48 (Cyto) | 0.022 (Cyto) | 0.81 (Cyto) | 0.95 (Nuc) | 0.003 (Nuc) | 0.352 (Nuc) |
| PSEN1 | 0.66 (Nuc); 1.3 (Cyto) | 0.007 (Nuc); $8.85 \times 10^{-5}$ (Cyto) | 0.825 (Nuc); **0.001 (Cyto)** | | | | 0.50 (Nuc); 0.95 (Cyto) | 0.023 (Nuc); 0.001 (Cyto) | 0.651 (Nuc); **0.041 (Cyto)** |
| EDC4 | -1.7 (Nuc); -1.06 (Cyto) | 0.025 (Nuc); $4.74 \times 10^{-5}$ (Cyto) | 0.91 (Nuc); **0.0007 (Cyto)** | -0.58 (Cyto) | 0.004 (Cyto) | 0.402 (Cyto) | -0.48 (Cyto) | 0.017 (Cyto) | 0.401 (Cyto) |
| MAP1LC3B | -3.03 (Cyto) | 0.0002 (Cyto) | **0.002 (Cyto)** | -2.88 (Cyto) | 0.0001 (Cyto) | **0.008 (Cyto)** | | | |
| PTMS | 4.38 (Nuc) | 0.009 (Nuc) | 0.846 (Nuc) | 4.46 (Nuc); | 0.006 (Nuc); 1.48 × | 0.623 (Nuc); | -4.26 (Cyto) | $3.22 \times 10^{-6}$ (Cyto) | **$3.20 \times 10^{-8}$ (Cyto)** |

| | | | | | | | | | |
|---|---|---|---|---|---|---|---|---|---|
| | | | | 4.27 (Cyto) | $10^{-6}$ (Cyto) | **7.25 × $10^{-8}$ (Cyto)** | | | |
| PTMA | | | | 2.63 (Nuc); 2.1 (Cyto) | 0.008 (Nuc); 0.0003 (Cyto) | 0.654 (Nuc); **0.034 (Cyto)** | -2.64 (Cyto) | 5.31 × $10^{-5}$ (Cyto) | **0.0002 (Cyto)** |
| HOOK3 | | | | 0.610 (Cyto) | 0.031 (Cyto) | 0.862 (Cyto) | -1.16 (Cyto) | 0.0007 (Cyto) | **0.027 (Cyto)** |
| GEMIN5 | 3.64 (Cyto) | 0.0001 (Cyto) | **0.002 (Cyto)** | 2.03 (Cyto) | 0.007 (Cyto) | 0.512 (Cyto) | 1.62 (Cyto) | 0.032 (Cyto) | 0.524 (Cyto) |
| PSMC6 | -1.05 (Cyto) | 0.0002 (Cyto) | **0.002 (Cyto)** | | | | -0.68 (Cyto) | 0.004 (Cyto) | 0.149 (Cyto) |
| DYNLRB1 | 1.79 (Nuc); 1.59 (Cyto) | 0.0007 (Nuc); 0.009 (Cyto) | 0.497 (Nuc); 0.249 (Cyto) | 1.50 (Nuc); 2.20 (Cyto) | 0.002 (Nuc); 0.0004 (Cyto) | 0.479 (Nuc); 0.061 (Cyto) | | | |
| NCBP1 | -0.54 (Nuc); -0.55 (Cyto) | 0.026 (Nuc); 0.016 (Cyto) | 0.91 (Nuc); 0.337 (Cyto) | 0.44 (Cyto) | 0.028 (Cyto) | 0.847 (Cyto) | -0.99 (Cyto) | 0.0001 (Cyto) | **0.001 (Cyto)** |
| ANP32A | 1.88 (Nuc); -0.75 (Cyto) | 0.016 (Nuc); 0.020 (Cyto) | 0.888 (Nuc); 0.376 (Cyto) | 2.58 (Nuc) | 0.001 (Nuc) | 0.435 (Nuc) | -1.29 (Cyto) | 0.0003 (Cyto) | **0.007 (Cyto)** |
| GLIPR2 | 1.79 (Cyto) | 0.0002 (Cyto) | **0.003 (Cyto)** | | | | 1.41 (Cyto) | 0.001 (Cyto) | **0.045 (Cyto)** |
| TRNT1 | 2.26 (Cyto) | 0.0003 (Cyto) | **0.005 (Cyto)** | 2.25 (Cyto) | 0.0001 (Cyto) | **0.009 (Cyto)** | | | |
| CLTA | 0.87 (Cyto) | 0.0005 (Cyto) | **0.012 (Cyto)** | 0.45 (Cyto) | 0.024 (Cyto) | 0.824 (Cyto) | 0.42 (Cyto) | 0.042 (Cyto) | 0.587 (Cyto) |
| TAX1BP3 | 2.68 (Cyto) | 0.0005 (Cyto) | **0.013 (Cyto)** | | | | 3.3 (Cyto) | 5.30 × $10^{-5}$ (Cyto) | **0.0002 (Cyto)** |
| PI4KA | 1.24 (Cyto) | 0.0006 (Cyto) | **0.014 (Cyto)** | | | | 0.87 (Cyto) | 0.006 (Cyto) | 0.206 (Cyto) |
| CLMP | 1.8 (Cyto) | 0.0006 (Cyto) | **0.014 (Cyto)** | 1.18 (Cyto) | 0.006 (Cyto) | 0.506 (Cyto) | | | |
| THOC5 | 1.11 (Cyto) | 0.0008 (Cyto) | **0.021 (Cyto)** | | | | 1.28 (Cyto) | 0.0002 (Cyto) | **0.001 (Cyto)** |
| EXOC6B | 1.36 (Cyto) | 0.0009 (Cyto) | **0.024 (Cyto)** | | | | 0.99 (Cyto) | 0.007 (Cyto) | 0.221 (Cyto) |
| DNPEP | 3.30 (Cyto) | 0.001 (Cyto) | **0.033 (Cyto)** | | | | 3.37 (Cyto) | 0.0006 (Cyto) | **0.023 (Cyto)** |
| GOLGA6L6 | -1.95 (Cyto) | 0.001 (Cyto) | **0.047 (Cyto)** | | | | -1.5 (Cyto) | 0.007 (Cyto) | 0.225 (Cyto) |

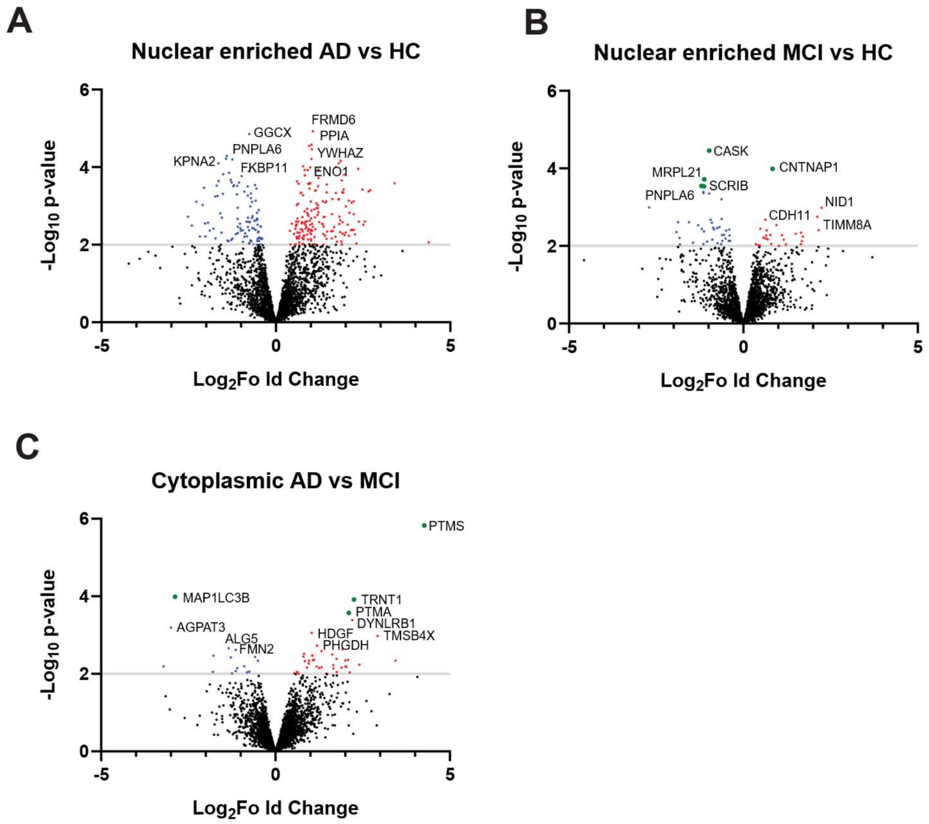


**Fig. S2. Additional volcano plots for fraction-specific proteomic comparisons.** Volcano plots showing differentially abundant proteins in (**A**) the nuclear-enriched AD vs HC, (**B**) the nuclear-enriched MCI vs HC and (**C**) the cytoplasmic AD vs MCI comparison. Green points indicate proteins that remained significant after FDR correction. Red and blue points indicate proteins meeting the nominal significance threshold of $p < 0.01$, with red representing increased abundance and blue representing decreased abundance in the first group of each comparison. The horizontal line indicates $p = 0.01$. Pairwise statistical analysis was performed using Bayesian statistical testing (LFQ-Analyst).

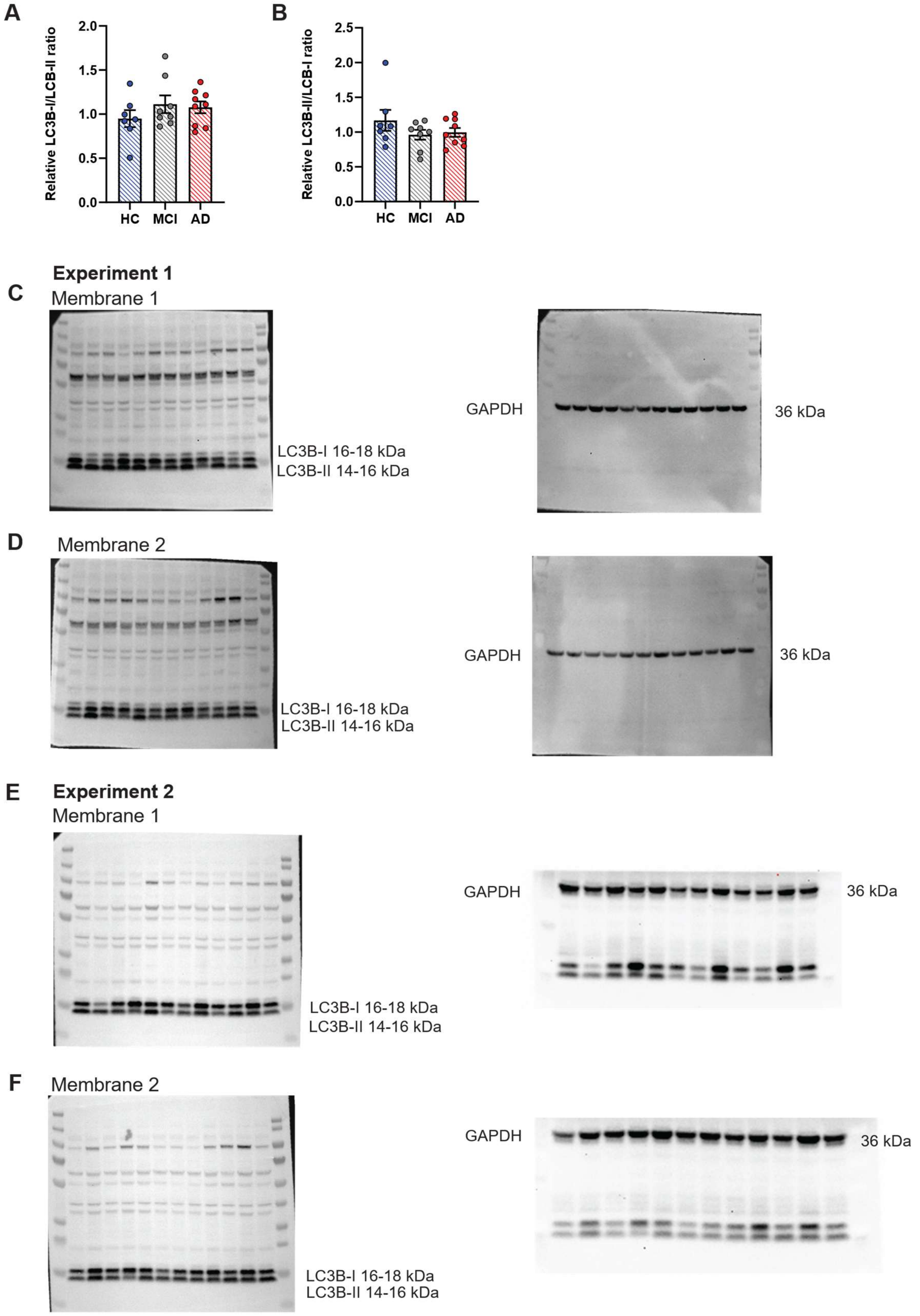


**Fig. S3. Additional LC3B western blot quantification and full membranes. (A)** Relative LC3B-I/LC3B-II ratio and (**B**) relative LC3B-II/LC3B-I ratio in HC (n = 7), MCI (n = 8) and AD (n = 9) ONS cells. Ratios were calculated from LC3B-I and LC3B-II band intensities obtained across two independent western blot experiments.

No significant group-level differences were detected. **(C, D)** Full LC3B membranes and corresponding GAPDH membranes from experiment 1. **(E, F)** Full LC3B membranes and corresponding GAPDH membranes from experiment 2. Sample order is the same as in Figure 3K. Data are shown as mean ± SEM.

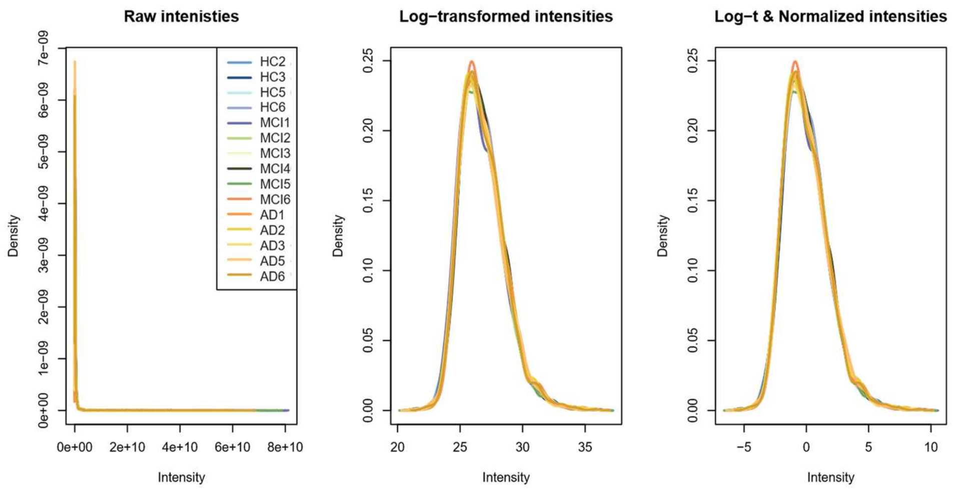


**Fig. S4. Quality control of reconstructed bulk proteomics data.** Density plots showing peptide intensity distributions before and after data processing. Raw intensities are shown before transformation, followed by log-transformed intensities and log-transformed normalised intensities across samples. The similar distribution of log-transformed normalised intensities indicates comparable intensity distributions across samples after processing.

**Table S2. Proteins significantly altered in AD ONS cells compared with HC in the reconstructed bulk proteomics dataset.** The reconstructed bulk proteomics dataset identified 40 differentially abundant proteins in AD ONS cells compared with HC after FDR correction, using an adjusted $p$ < 0.05 threshold. $Log_2FC$, average expression, nominal $p$-values and Benjamini-Hochberg adjusted $p$-values are shown.

| | Protein | $Log_2FC$ | Average expression | $p$-value | Adjusted $p$-value |
|---|---|---|---|---|---|
| 1 | ALG1 | -1.67041 | -1.46313 | 3.30E-07 | 0.000822 |
| 2 | AGRN | -1.66844 | -1.25114 | 1.41E-05 | 0.01507 |
| 3 | POSTN | -1.58274 | 2.414809 | 2.31E-05 | 0.01507 |
| 4 | FKBP11 | -1.49874 | 2.357372 | 2.55E-05 | 0.01507 |
| 5 | SLC4A7 | -1.08545 | -0.84795 | 3.03E-05 | 0.01507 |
| 6 | BCL2L13 | -1.62689 | -0.91782 | 4.52E-05 | 0.01874 |
| 7 | CTNNA2 | -1.9679 | 0.980095 | 6.03E-05 | 0.020618 |
| 8 | SEC11A | -0.92815 | 0.791566 | 7.24E-05 | 0.020618 |
| 9 | S100A4 | 1.592891 | -0.59303 | 7.77E-05 | 0.020618 |
| 10 | ARHGEF7 | -0.98624 | -1.99213 | 8.28E-05 | 0.020618 |
| 11 | SUSD2 | -4.02781 | -1.13172 | 9.30E-05 | 0.02106 |

| | | | | | |
|---|---|---|---|---|---|
| 12 | ATP2B1 | -0.98562 | -1.70589 | 0.000123 | 0.025524 |
| 13 | NOL11 | -0.86483 | -1.03221 | 0.000138 | 0.026391 |
| 14 | MAK16 | -0.8617 | -0.83696 | 0.000167 | 0.028415 |
| 15 | LAMTOR5 | 2.145765 | -0.78591 | 0.000171 | 0.028415 |
| 16 | SYNE1 | -0.70856 | 4.286791 | 0.000205 | 0.031804 |
| 17 | GOLT1B | -1.32586 | -1.51582 | 0.000219 | 0.031804 |
| 18 | PRXL2A | -1.18475 | 0.781841 | 0.000233 | 0.031804 |
| 19 | STMN1 | 1.498883 | 0.249366 | 0.000248 | 0.031804 |
| 20 | SIL1 | -1.09755 | -1.69617 | 0.000276 | 0.031804 |
| 21 | PSAP | 1.400715 | 1.229318 | 0.000305 | 0.031804 |
| 22 | DHCR24 | 0.987651 | -1.27144 | 0.000312 | 0.031804 |
| 23 | MRPL21 | -1.1533 | -0.85868 | 0.000315 | 0.031804 |
| 24 | CNTNAP1 | 0.912595 | -1.72886 | 0.000332 | 0.031804 |
| 25 | RALB | 0.798678 | -1.10919 | 0.000344 | 0.031804 |
| 26 | EDC4 | -0.80497 | -1.06473 | 0.000359 | 0.031804 |
| 27 | DYNLRB1 | 1.386328 | -1.5694 | 0.000364 | 0.031804 |
| 28 | CLTA | 0.857551 | 1.133547 | 0.00037 | 0.031804 |
| 29 | ARSB | -1.20478 | -0.7093 | 0.00037 | 0.031804 |
| 30 | LRRC59 | -0.80195 | 5.220553 | 0.000398 | 0.033009 |
| 31 | KRR1 | -0.98687 | 0.698744 | 0.000444 | 0.034677 |
| 32 | TOP2A | -1.0361 | -1.11378 | 0.000446 | 0.034677 |
| 33 | MRPL9 | -0.82862 | -1.34954 | 0.000493 | 0.037123 |
| 34 | OCIAD2 | -1.55765 | -0.99298 | 0.000507 | 0.037123 |
| 35 | NPLOC4 | -0.90849 | -1.07307 | 0.000569 | 0.039836 |
| 36 | TMSB4X | 1.967886 | 0.517748 | 0.000576 | 0.039836 |
| 37 | PUS7 | -1.27159 | -0.07015 | 0.000595 | 0.040024 |
| 38 | BASP1 | 1.52084 | 3.31865 | 0.000644 | 0.042167 |
| 39 | OCC1 | 0.756804 | 0.604816 | 0.000674 | 0.043042 |
| 40 | COPS7A | 0.951403 | -1.58689 | 0.000697 | 0.043385 |

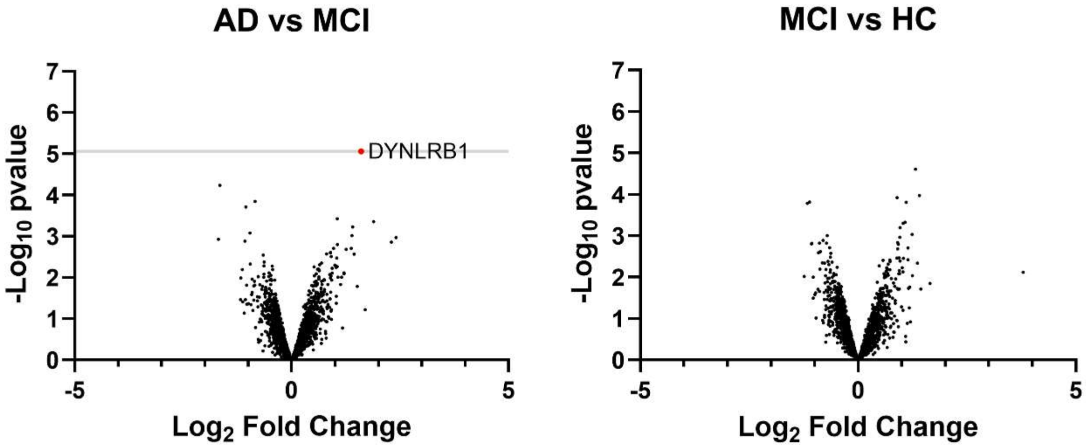


**Fig. S5. Differential protein abundance in reconstructed bulk proteomics for AD vs MCI and MCI vs HC comparisons.** Volcano plots showing differential protein abundance in (**A**) AD vs MCI and (**B**) MCI vs HC reconstructed bulk proteomics comparisons. The x-axis shows $\log_2$ fold change and the y-axis shows $-\log_{10}$-transformed nominal *p*-values. Proteins remaining significant after FDR correction (adjusted $p < 0.05$) are highlighted. DYNLRB1 was the only protein that remained significant after FDR correction in the AD vs MCI comparison, whereas no proteins remained significant after FDR correction in the MCI vs HC comparison.

### Experiment 1

Membrane 1

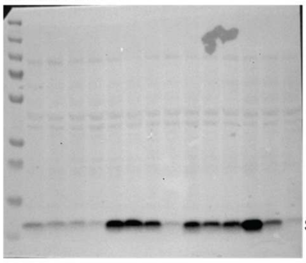

S100A4 12 kDa

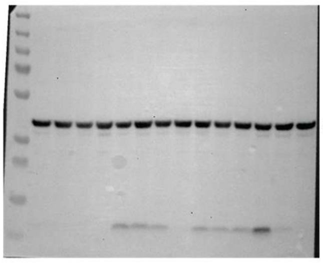

GAPDH 36 kDa

Membrane 2

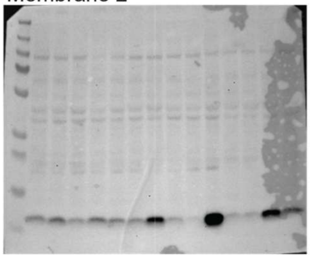

S100A4 12 kDa

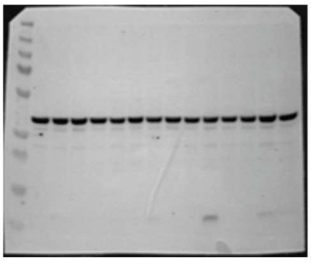

GAPDH 36 kDa

### Experiment 2

Membrane 1

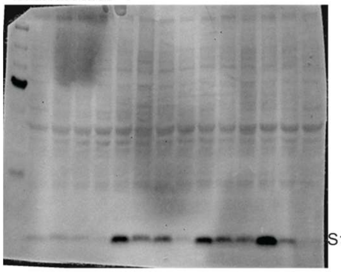

S100A4 12 kDa

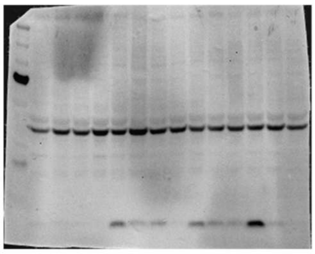

GAPDH 36 kDa

Membrane 2

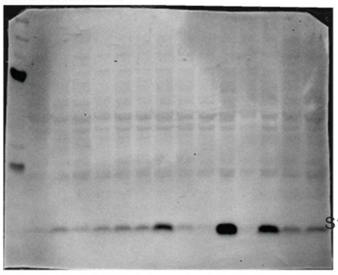

S100A4 12 kDa

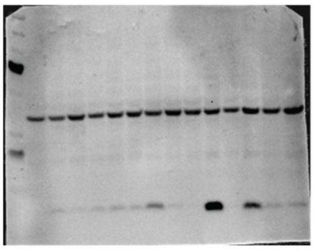

GAPDH 36 kDa

### Experiment 3

Membrane 1

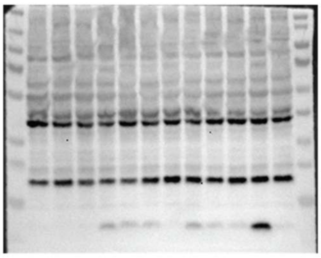

GAPDH 36 kDa

S100A4 12 kDa

Membrane 2

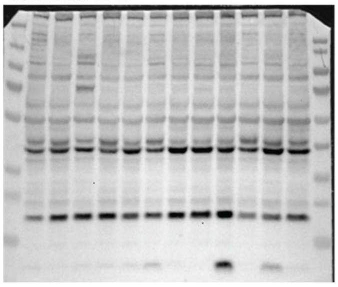

GAPDH 36 kDa

S100A4 12 kDa

**Fig. S6. Full western blot membranes for S100A4 validation.** Full western blot membranes showing S100A4 and corresponding GAPDH loading controls from three independent experiments. Experiments 1 and 2 were performed across two membranes, while experiment 3 was performed across two membranes with S100A4 and GAPDH detected on the same blots. The expected molecular weights for S100A4 (~12 kDa) and GAPDH (~36 kDa) are indicated. The sample order is the same as in Figure 5F.

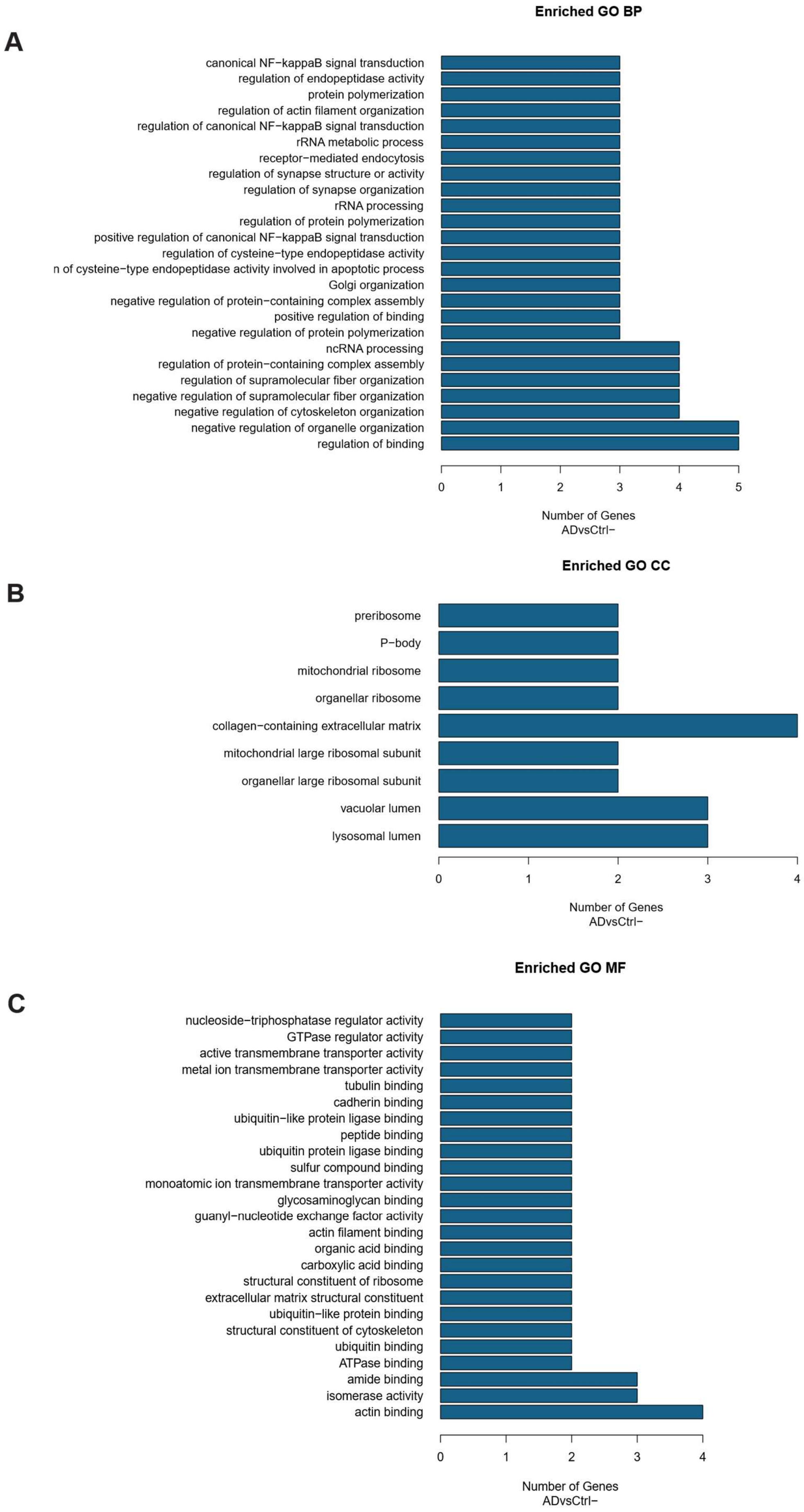
A
Enriched GO BP
canonical NF−kappaB signal transduction
regulation of endopeptidase activity
protein polymerization
regulation of actin filament organization
regulation of canonical NF−kappaB signal transduction
rRNA metabolic process
receptor−mediated endocytosis
regulation of synapse structure or activity
regulation of synapse organization
rRNA processing
regulation of protein polymerization
positive regulation of canonical NF−kappaB signal transduction
regulation of cysteine−type endopeptidase activity
n of cysteine−type endopeptidase activity involved in apoptotic process
Golgi organization
negative regulation of protein−containing complex assembly
positive regulation of binding
negative regulation of protein polymerization
ncRNA processing
regulation of protein−containing complex assembly
regulation of supramolecular fiber organization
negative regulation of supramolecular fiber organization
negative regulation of cytoskeleton organization
negative regulation of organelle organization
regulation of binding
0 1 2 3 4 5
Number of Genes
ADvsCtrl−
B
Enriched GO CC
preribosome
P−body
mitochondrial ribosome
organellar ribosome
collagen−containing extracellular matrix
mitochondrial large ribosomal subunit
organellar large ribosomal subunit
vacuolar lumen
lysosomal lumen
0 1 2 3 4
Number of Genes
ADvsCtrl−
C
Enriched GO MF
nucleoside−triphosphatase regulator activity
GTPase regulator activity
active transmembrane transporter activity
metal ion transmembrane transporter activity
tubulin binding
cadherin binding
ubiquitin−like protein ligase binding
peptide binding
ubiquitin protein ligase binding
sulfur compound binding
monoatomic ion transmembrane transporter activity
glycosaminoglycan binding
guanyl−nucleotide exchange factor activity
actin filament binding
organic acid binding
carboxylic acid binding
structural constituent of ribosome
extracellular matrix structural constituent
ubiquitin−like protein binding
structural constituent of cytoskeleton
ubiquitin binding
ATPase binding
amide binding
isomerase activity
actin binding
0 1 2 3 4
Number of Genes
ADvsCtrl−

**Fig. S7. Gene Ontology enrichment analysis of AD vs HC reconstructed bulk proteomics DEPs.** GO enrichment analysis was performed on the 40 FDR-significant AD vs HC reconstructed bulk proteomics DEPs using ClusterProfiler. **(A)** Full enriched GO biological process (BP) terms. **(B)** Enriched cellular component (CC) terms. **(C)** Enriched molecular function (MF) terms. The x-axis indicates the number of DEPs contributing to each enriched term.

**Table S3. Statistically significant ($p$ < 0.05) RNA-protein correlations between transcriptomic and proteomic data.**

| Name | shapiro_rna_pval | shapiro_prot_pval | cor_prot_rna | cor_pval | cor_pval_CI1 | cor_pval_CI2 |
|---|---|---|---|---|---|---|
| ABCC1 | 0.9232871 | 0.2121862 | -0.5181781 | 0.0478416 | -0.8142924 | -0.0080529 |
| AFAP1 | 0.4857471 | 0.2377327 | 0.7063804 | 0.0032433 | 0.3041891 | 0.8948424 |
| AHNAK | 0.8495898 | 0.5758205 | 0.717126 | 0.0026198 | 0.3238211 | 0.8990987 |
| ALDH18A1 | 0.9547105 | 0.6020168 | -0.6784912 | 0.0054242 | -0.8836333 | -0.2547817 |
| ALDH9A1 | 0.5171519 | 0.3949811 | 0.5217456 | 0.0460632 | 0.0129417 | 0.8159333 |
| ANPEP | 0.336126 | 0.1608808 | 0.8677209 | 0.0000277 | 0.6398899 | 0.9553359 |
| ANXA1 | 0.5506715 | 0.2874992 | 0.698697 | 0.0037576 | 0.2903582 | 0.8917779 |
| ASPH | 0.0582086 | 0.4383643 | 0.5239449 | 0.0449915 | 0.0159678 | 0.8169424 |
| CAVIN2 | 0.7667424 | 0.338604 | 0.6007357 | 0.0178724 | 0.1278019 | 0.851089 |
| CBR1 | 0.7600115 | 0.4440199 | 0.6921245 | 0.0042472 | 0.2786607 | 0.8891422 |
| CERCAM | 0.5377741 | 0.7024489 | 0.5872711 | 0.0213441 | 0.1072829 | 0.845251 |
| CMAS | 0.8032619 | 0.9880514 | 0.5311838 | 0.041594 | 0.0259941 | 0.8202515 |
| COL12A1 | 0.3679279 | 0.4060158 | 0.5521138 | 0.0328404 | 0.0555668 | 0.8297115 |
| COL16A1 | 0.3510363 | 0.4756792 | 0.6866197 | 0.0046951 | 0.268957 | 0.8869247 |
| COL18A1 | 0.0930615 | 0.6760156 | 0.8010233 | 0.0003319 | 0.4897021 | 0.9311877 |
| COL4A1 | 0.0920643 | 0.7776718 | 0.5994983 | 0.018172 | 0.1258992 | 0.8505551 |
| COL5A2 | 0.4087674 | 0.9710158 | 0.547532 | 0.034626 | 0.0490176 | 0.8276542 |
| COL6A1 | 0.4245123 | 0.0715025 | 0.8321867 | 0.0001188 | 0.5576485 | 0.9426146 |
| COL6A3 | 0.8440396 | 0.0996993 | 0.6175419 | 0.0141672 | 0.1539942 | 0.85829 |
| COPB1 | 0.0985443 | 0.7054351 | 0.6671518 | 0.0065876 | 0.2353066 | 0.8790076 |
| CORO1B | 0.632154 | 0.7860495 | -0.5296784 | 0.0422843 | -0.819565 | -0.0239007 |
| CORO2B | 0.7351485 | 0.6788842 | 0.7651869 | 0.0008871 | 0.4159776 | 0.9177234 |
| CRYZ | 0.2211299 | 0.9128272 | 0.7831207 | 0.0005547 | 0.4523069 | 0.9245055 |
| CUL4B | 0.245183 | 0.8617668 | -0.6230821 | 0.0130863 | -0.8606433 | -0.1627737 |
| CYB5A | 0.8261026 | 0.2367603 | 0.6643364 | 0.0069049 | 0.2305244 | 0.8778529 |
| CYB5B | 0.775616 | 0.835908 | 0.5515196 | 0.033068 | 0.054715 | 0.8294451 |
| CYC1 | 0.2658208 | 0.6477995 | 0.5553069 | 0.0316375 | 0.0601564 | 0.8311409 |
| DDX17 | 0.6099264 | 0.0909486 | 0.5233295 | 0.0452895 | 0.0151201 | 0.8166602 |
| DNAJB4 | 0.2765468 | 0.4171355 | 0.5941558 | 0.0195101 | 0.1177238 | 0.8482438 |
| DNM2 | 0.0526006 | 0.3013226 | 0.6227095 | 0.013157 | 0.1621809 | 0.8604853 |
| DPY30 | 0.3268908 | 0.9353735 | -0.683988 | 0.0049221 | -0.8858613 | -0.2643477 |

| | | | | | | |
|---|---|---|---|---|---|---|
| DSP | 0.2065384 | 0.189598 | 0.522436 | 0.0457248 | 0.0138906 | 0.8162502 |
| DYNC1I2 | 0.6404359 | 0.4237337 | 0.5905744 | 0.0204484 | 0.1122793 | 0.846689 |
| DYNLL1 | 0.1110732 | 0.6007838 | 0.6425675 | 0.0097836 | 0.194242 | 0.86884 |
| EDIL3 | 0.9909385 | 0.9971106 | 0.6152003 | 0.0146443 | 0.1503053 | 0.8572924 |
| EHD4 | 0.3615238 | 0.8454806 | 0.6726043 | 0.0060059 | 0.2446281 | 0.8812368 |
| ELOVL1 | 0.7731241 | 0.0524529 | -0.5924389 | 0.0199557 | -0.847499 | -0.1151102 |
| ENDOD1 | 0.2507519 | 0.659522 | 0.6071668 | 0.0163759 | 0.1377473 | 0.8538557 |
| EPHA2 | 0.8297995 | 0.7990859 | 0.6666903 | 0.0066388 | 0.2345213 | 0.8788185 |
| FADS2 | 0.0699948 | 0.8638967 | 0.5516545 | 0.0330162 | 0.0549084 | 0.8295056 |
| FAM3C | 0.4579258 | 0.6011617 | 0.5269548 | 0.0435547 | 0.0201243 | 0.8183207 |
| FLNB | 0.9514409 | 0.2125432 | 0.5283762 | 0.0428882 | 0.0220933 | 0.8189704 |
| FLNC | 0.8605308 | 0.8987371 | 0.6088225 | 0.0160067 | 0.1403233 | 0.8545658 |
| FZD7 | 0.6607313 | 0.2189366 | 0.5271759 | 0.0434506 | 0.0204303 | 0.8184218 |
| GANAB | 0.7568411 | 0.7095216 | -0.5349494 | 0.0399039 | -0.8219652 | -0.0312502 |
| GAPDH | 0.9532214 | 0.6367467 | 0.5707403 | 0.0262847 | 0.0826384 | 0.8379984 |
| GDI2 | 0.7294632 | 0.999098 | 0.5768051 | 0.0243809 | 0.0916112 | 0.8406702 |
| GGH | 0.5190312 | 0.5954266 | 0.7768067 | 0.0006575 | 0.4393901 | 0.9221279 |
| GLIPR2 | 0.4105917 | 0.7850838 | 0.7616397 | 0.0009688 | 0.4089216 | 0.9163714 |
| GLS | 0.0618202 | 0.4153423 | 0.6078197 | 0.0162296 | 0.1387624 | 0.8541359 |
| GNB4 | 0.8106636 | 0.8317408 | 0.5855388 | 0.0218257 | 0.1046724 | 0.8444954 |
| GPX7 | 0.0667754 | 0.5473446 | 0.5870849 | 0.0213955 | 0.107002 | 0.8451699 |
| GRAMD2B | 0.7997625 | 0.8365156 | 0.6741855 | 0.005845 | 0.2473462 | 0.8818816 |
| GREM1 | 0.1637385 | 0.7504302 | 0.5546573 | 0.0318795 | 0.059221 | 0.8308504 |
| GULP1 | 0.2158821 | 0.2263758 | 0.6154245 | 0.0145981 | 0.1506579 | 0.857388 |
| HK2 | 0.7930973 | 0.350603 | 0.5289692 | 0.0426124 | 0.0229159 | 0.8192412 |
| HM13 | 0.4335455 | 0.1666938 | -0.6173924 | 0.0141973 | -0.8582264 | -0.1537582 |
| HSD17B12 | 0.6905391 | 0.1120524 | 0.6705547 | 0.0062196 | 0.2411149 | 0.8803999 |
| IDH2 | 0.212942 | 0.5815702 | 0.5767057 | 0.0244112 | 0.0914635 | 0.8406265 |
| IGFBP7 | 0.2688902 | 0.9423444 | 0.5845158 | 0.0221139 | 0.103134 | 0.8440488 |
| KCNMA1 | 0.6021301 | 0.9964815 | 0.6946539 | 0.0040531 | 0.283148 | 0.8901581 |
| LAP3 | 0.9917017 | 0.4184754 | 0.5693603 | 0.0267333 | 0.0806076 | 0.8373886 |
| LBR | 0.9425692 | 0.9665803 | 0.5261109 | 0.0439541 | 0.0189572 | 0.8179346 |
| LIPA | 0.1599155 | 0.9904026 | 0.5147251 | 0.0496107 | 0.0033439 | 0.8126997 |
| LNPK | 0.4457325 | 0.7479872 | 0.6199932 | 0.0136808 | 0.1578696 | 0.8593325 |
| LRPPRC | 0.8794804 | 0.0531062 | 0.5369018 | 0.039048 | 0.0339865 | 0.8228518 |
| LUZP1 | 0.4722012 | 0.6835997 | 0.6731142 | 0.0059536 | 0.2455038 | 0.8814448 |
| MANF | 0.3526329 | 0.3963366 | -0.5201485 | 0.0468532 | -0.8151992 | -0.01075 |
| MAPK1 | 0.8418577 | 0.2383172 | 0.5752774 | 0.0248503 | 0.0893435 | 0.8399984 |
| MFGE8 | 0.1933809 | 0.935157 | 0.6573626 | 0.0077426 | 0.2187684 | 0.8749819 |
| MXRA8 | 0.4623048 | 0.0772157 | 0.6033654 | 0.0172483 | 0.1318572 | 0.852222 |
| MYO1B | 0.1386059 | 0.9891914 | 0.808884 | 0.0002607 | 0.5064916 | 0.9340944 |
| NDUFS2 | 0.9567502 | 0.9957506 | 0.5157185 | 0.0490969 | 0.0046964 | 0.8131584 |

| | | | | | | |
|---|---|---|---|---|---|---|
| NDUFV1 | 0.9991242 | 0.9892016 | 0.7606348 | 0.0009931 | 0.4069303 | 0.9159877 |
| NES | 0.0977712 | 0.820958 | 0.8059702 | 0.0002854 | 0.5002413 | 0.9330189 |
| NGDN | 0.2536874 | 0.8765961 | -0.6112235 | 0.0154828 | -0.8555939 | -0.1440701 |
| NUMA1 | 0.9499559 | 0.4154667 | 0.5436724 | 0.0361857 | 0.0435339 | 0.8259153 |
| NUP35 | 0.4526461 | 0.9315071 | 0.5626369 | 0.0290018 | 0.0707717 | 0.8344083 |
| NUP37 | 0.4376771 | 0.2788451 | 0.5828965 | 0.0225761 | 0.1007032 | 0.8433409 |
| OGDH | 0.1461844 | 0.7619016 | 0.6449127 | 0.0094348 | 0.1980927 | 0.8698182 |
| PAK2 | 0.9676866 | 0.8221911 | 0.5708915 | 0.0262359 | 0.0828611 | 0.8380651 |
| PAPSS1 | 0.8687901 | 0.3639659 | 0.5886657 | 0.0209624 | 0.1093894 | 0.8458586 |
| PFN2 | 0.1590168 | 0.1596164 | 0.7286072 | 0.0020636 | 0.345178 | 0.9036086 |
| PKM | 0.1726183 | 0.3992139 | 0.5322025 | 0.0411316 | 0.0274132 | 0.8207156 |
| PLOD3 | 0.1987657 | 0.5490463 | -0.552007 | 0.0328812 | -0.8296637 | -0.0554137 |
| PPP2R2A | 0.2904188 | 0.7584508 | 0.5212351 | 0.0463147 | 0.0122406 | 0.8156987 |
| PVR | 0.2262783 | 0.6712296 | 0.8796745 | 0.0000154 | 0.6687806 | 0.9595435 |
| PXN | 0.2709461 | 0.3415215 | 0.6225435 | 0.0131885 | 0.161917 | 0.860415 |
| RAB23 | 0.144517 | 0.9998276 | 0.5861455 | 0.0216561 | 0.105586 | 0.8447602 |
| RAB3B | 0.7936553 | 0.1738131 | 0.6380941 | 0.0104767 | 0.1869351 | 0.8669691 |
| RAB8B | 0.6630518 | 0.6695563 | 0.5655172 | 0.0280129 | 0.0749738 | 0.835687 |
| RAP1B | 0.5297258 | 0.8189814 | 0.6073326 | 0.0163387 | 0.138005 | 0.8539269 |
| RASA3 | 0.1037729 | 0.178311 | -0.5436594 | 0.0361911 | -0.8259094 | -0.0435154 |
| RBM3 | 0.37419 | 0.9820599 | 0.6859275 | 0.004754 | 0.2677428 | 0.8866452 |
| RBM42 | 0.9928836 | 0.9057857 | 0.7304335 | 0.0019847 | 0.3486123 | 0.9043225 |
| RDH14 | 0.5243894 | 0.7284682 | 0.62375 | 0.0129605 | 0.163837 | 0.8609263 |
| RECK | 0.3321959 | 0.293808 | 0.7666882 | 0.0008542 | 0.4189767 | 0.9182946 |
| RND3 | 0.4948545 | 0.4050077 | 0.611847 | 0.015349 | 0.1450451 | 0.8558605 |
| RPA2 | 0.0549468 | 0.6562882 | -0.518321 | 0.0477694 | -0.8143582 | -0.0082482 |
| RPLP1 | 0.9627111 | 0.2322886 | 0.672042 | 0.0060639 | 0.2436632 | 0.8810074 |
| RPP30 | 0.6454969 | 0.7999842 | 0.5728061 | 0.025624 | 0.0856858 | 0.8389099 |
| RRAS2 | 0.0651344 | 0.6569339 | 0.7032058 | 0.0034485 | 0.2984539 | 0.8935783 |
| SDCBP | 0.4481799 | 0.8992737 | 0.7704022 | 0.0007771 | 0.4264289 | 0.919705 |
| SEC61A1 | 0.8314991 | 0.5773041 | -0.7446307 | 0.0014499 | -0.9098387 | -0.3756635 |
| SEL1L | 0.9977886 | 0.9413897 | -0.7114921 | 0.0029334 | -0.8968714 | -0.3134856 |
| SERPINE1 | 0.7152183 | 0.6666569 | 0.7844441 | 0.0005349 | 0.455032 | 0.9250025 |
| SH3BGRL3 | 0.564971 | 0.1550647 | 0.649223 | 0.0088191 | 0.2052061 | 0.8716115 |
| SNX9 | 0.7451889 | 0.9883333 | 0.6434052 | 0.0096578 | 0.1956159 | 0.8691896 |
| SPARC | 0.9520206 | 0.224838 | 0.6479534 | 0.0089971 | 0.203106 | 0.8710839 |
| SRM | 0.3560032 | 0.6693148 | 0.653499 | 0.0082399 | 0.2123096 | 0.8733847 |
| SULF1 | 0.6349468 | 0.3506509 | 0.5964076 | 0.0189372 | 0.1211618 | 0.8492192 |
| SUSD2 | 0.8174643 | 0.2931193 | 0.6380039 | 0.0104911 | 0.1867882 | 0.8669313 |
| THBS1 | 0.0599797 | 0.8553332 | 0.6287252 | 0.0120522 | 0.1717918 | 0.8630298 |
| UCHL1 | 0.3885838 | 0.7956428 | 0.6495409 | 0.008775 | 0.2057326 | 0.8717436 |
| VCAM1 | 0.9888417 | 0.1252547 | 0.6615475 | 0.0072309 | 0.2258078 | 0.8767066 |

| | | | | | | |
|---|---|---|---|---|---|---|
| VDAC1 | 0.8971617 | 0.281614 | -0.6031381 | 0.0173016 | -0.8521242 | -0.1315061 |
| YWHAG | 0.815262 | 0.8377387 | 0.5972728 | 0.0187205 | 0.1224859 | 0.8495935 |

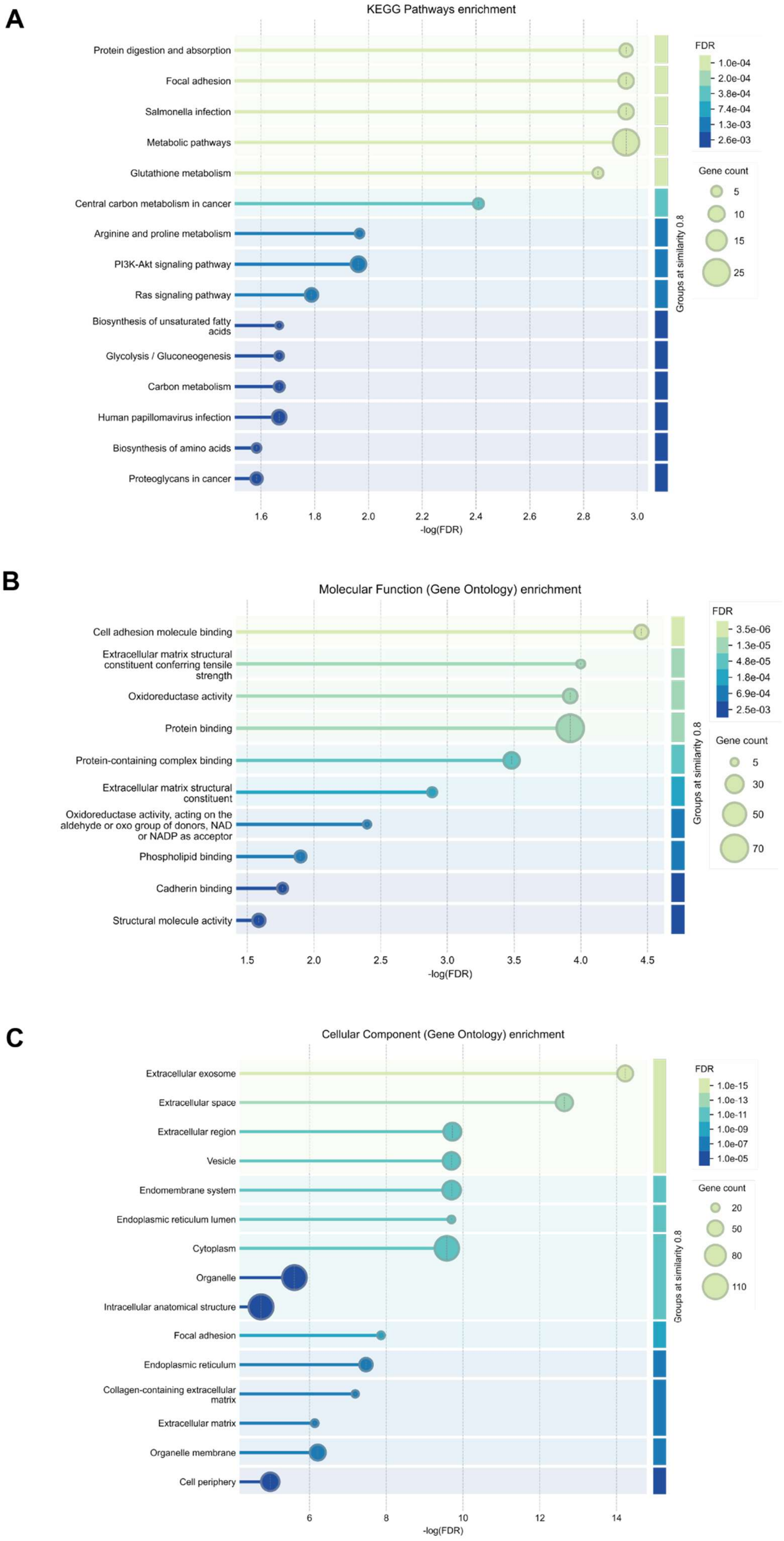

A
KEGG Pathways enrichment
Protein digestion and absorption
Focal adhesion
Salmonella infection
Metabolic pathways
Glutathione metabolism
Central carbon metabolism in cancer
Arginine and proline metabolism
PI3K-Akt signaling pathway
Ras signaling pathway
Biosynthesis of unsaturated fatty acids
Glycolysis / Gluconeogenesis
Carbon metabolism
Human papillomavirus infection
Biosynthesis of amino acids
Proteoglycans in cancer
1.6 1.8 2.0 2.2 2.4 2.6 2.8 3.0
-log(FDR)
Groups at similarity 0.8
FDR
1.0e-04
2.0e-04
3.8e-04
7.4e-04
1.3e-03
2.6e-03
Gene count
5
10
15
25
B
Molecular Function (Gene Ontology) enrichment
Cell adhesion molecule binding
Extracellular matrix structural constituent conferring tensile strength
Oxidoreductase activity
Protein binding
Protein-containing complex binding
Extracellular matrix structural constituent
Oxidoreductase activity, acting on the aldehyde or oxo group of donors, NAD or NADP as acceptor
Phospholipid binding
Cadherin binding
Structural molecule activity
1.5 2.0 2.5 3.0 3.5 4.0 4.5
-log(FDR)
Groups at similarity 0.8
FDR
3.5e-06
1.3e-05
4.8e-05
1.8e-04
6.9e-04
2.5e-03
Gene count
5
30
50
70
C
Cellular Component (Gene Ontology) enrichment
Extracellular exosome
Extracellular space
Extracellular region
Vesicle
Endomembrane system
Endoplasmic reticulum lumen
Cytoplasm
Organelle
Intracellular anatomical structure
Focal adhesion
Endoplasmic reticulum
Collagen-containing extracellular matrix
Extracellular matrix
Organelle membrane
Cell periphery
6 8 10 12 14
-log(FDR)
Groups at similarity 0.8
FDR
1.0e-15
1.0e-13
1.0e-11
1.0e-09
1.0e-07
1.0e-05
Gene count
20
50
80
110

**Fig. S8. Functional enrichment analysis of correlated genes.** Functional enrichment analysis was performed using genes showing significant RNA–protein correlations at $p < 0.05$. (**A**) KEGG pathway enrichment, (**B**) Gene Ontology (GO) molecular function enrichment and (**C**) GO cellular component enrichment. Only terms with FDR < 0.05 are shown. Bubble size indicates gene count, and colour indicates FDR-adjusted enrichment significance.

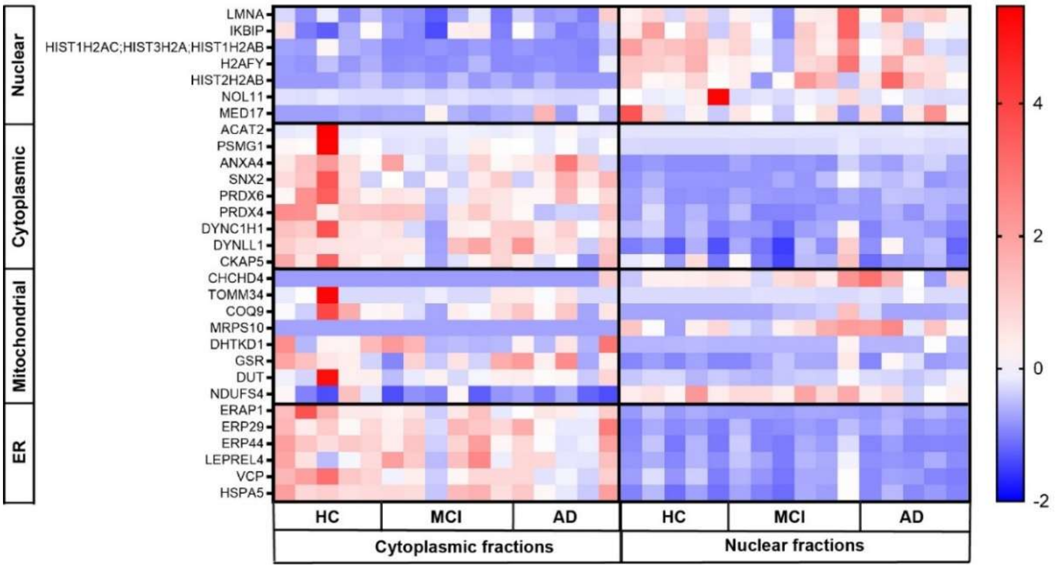


**Fig. S9. Separation of nuclear, cytoplasmic, mitochondrial and endoplasmic reticulum marker proteins across cytoplasmic and nuclear-enriched fractions.** Selected resident marker proteins for the nucleus ($n$=7), cytoplasm ($n$=9), mitochondria ($n$=8) and endoplasmic reticulum (ER) ($n$=6) were assessed across the cytoplasmic and nuclear-enriched fractions from HC ($n$=5), MCI ($n$=6) and AD ($n$=5) ONS cell lines. Data is shown as a Z-score normalised protein intensities to visualise the relative enrichment of compartment-associated marker proteins within each prepared fraction.

**Table S4. Summary of donor information**. Table adapted from (Rantanen et al., 2022).

| Study cohorts | | Healthy control (HC) | Mild cognitive impairment (MCI) | Alzheimer's disease (AD) |
|---|---|---|---|---|
| No. of participants | | $n = 4$ | $n = 6$ | $n = 5$ |
| Sex of participants | Females (%) | 50 % | 33.3 % | 60 % |
| | Males (%) | 50 % | 66.6 % | 40 % |
| Age of participants (mean ± SD[1]) | | 69 ± 4.2 | 72 ± 6.5 | 70 ± 9.0 |
| ApoE[2] genotype | E2/E3 (%) | 0 % | 16.6 % | 0 % |
| | E3/E3 (%) | 50 % | 50 % | 60 % |
| | E3/E4 (%) | 50 % | 33.3 % | 20.0 % |
| | E4/E4 (%) | 0 % | 0 % | 20.0 % |
| Olfactory function | Normal (%) | 50 % | 83.3 % | 20 % |
| | Hyposmia (%) | 50 % | 16.6 % | 40 % |
| | Anosmia (%) | 0 % | 0 % | 40 % |

[1.] SD = standard deviation; [2.] ApoE = apolipoprotein E.

**Table S5. Primer sequences used in the study.**

| Primer | Forward Sequence | Reverse Sequence |
|---|---|---|
| *TUBB3* | GGCCAAGTTCTGGGAAGTCAT | CTCGAGGCACGTACTTGTGA |
| *NESTIN* | CTCAGCTTTCAGGACCCCAA | GTCTCAAGGGTAGCAGGCAA |
| *PAX6* | TGGTATTCTCTCCCCCTCCT | TAAGGATGTTGAACGGGCAG |
| *S100B* | TTCTGGAAGGGAGGGAGACA | CTCCTGCTCTTTGATTTCCTCT |
| *MAP2* | GACTGCAGCTCTGCCTTTAG | AAGTAAATCTTCCTCCACTGTGA C |
| *18S* | TTCGAGGCCCTGTAATTGGA | GCAGCAACTTTAATATACGCTA TTGG |

**Table S6. Primary and secondary antibodies used for immunofluorescence characterisation of ONS cells.**

| **Primary antibodies** | **Species** | **Dilution** | **Company (Cat#)** |
|---|---|---|---|
| DCX | Rabbit | 1:200 | Cell Signalling Technology (4604S) |
| TUBB3 | Mouse | 1:500 | BioLegend (801202) |
| Nestin | Mouse | 1:200 | Abcam (ab22035) |
| AQP4 | Mouse | 1:200 | Abcam (ab9512) |
| S100B | Rabbit | 1:200 | Abcam, (ab52642) |
| **Secondary antibodies** | **Species** | **Dilution** | **Company (Cat#)** |
| Anti-Mouse Alexa Fluor 488 | Goat | 1:250 | Invitrogen (A-11029) |
| Anti-Rabbit Alexa Fluor 594 | Goat | 1:250 | Invitrogen (A-11037) |
| Anti-Rat Alexa Fluor 647 | Goat | 1:250 | Invitrogen (A-21247) |
